# RiboBA: a bias-aware probabilistic framework for robust ORF identification across diverse ribosome profiling protocols

**DOI:** 10.64898/2026.03.17.712439

**Authors:** Junyu Bai, Ruolin Yang

## Abstract

By mapping ribosome-protected fragments (RPFs) genome-wide, ribosome profiling (Ribo–seq) has uncovered extensive translation beyond conventional coding sequences, revealing non-canonical ORFs (ncORFs) with emerging roles in diverse biological processes. However, protocol-induced biases introduced during library construction can substantially distort RPF signals. Most existing ORF callers are not designed to explicitly account for such artifacts, limiting robust ncORF identification. Here, we present RiboBA, a bias-aware probabilistic framework to address this challenge. RiboBA consists of two main components: a generative module that recovers protocol-induced biases and codon-level ribosome occupancy, and a supervised module that identifies translated ORFs and initiation sites using the resulting bias-adjusted profiles. Evaluated through simulations and on a range of Ribo–seq datasets—particularly supported by cell-type-specific immunopeptidomics—RiboBA robustly recovers protocol-induced parameters and achieves superior accuracy and sensitivity in ncORF identification. Notably, RiboBA performs particularly well on RNase I libraries with attenuated three-nucleotide periodicity, as well as on MNase and nuclease P1 libraries, while maintaining competitive runtimes. In a Drosophila case study, RiboBA identifies conserved ncORFs with coding potential, including recurrent upstream translation of ThrRS and Mettl2 that suggests a potential threonine-specific translational control axis.

**Graphical Abstract:** 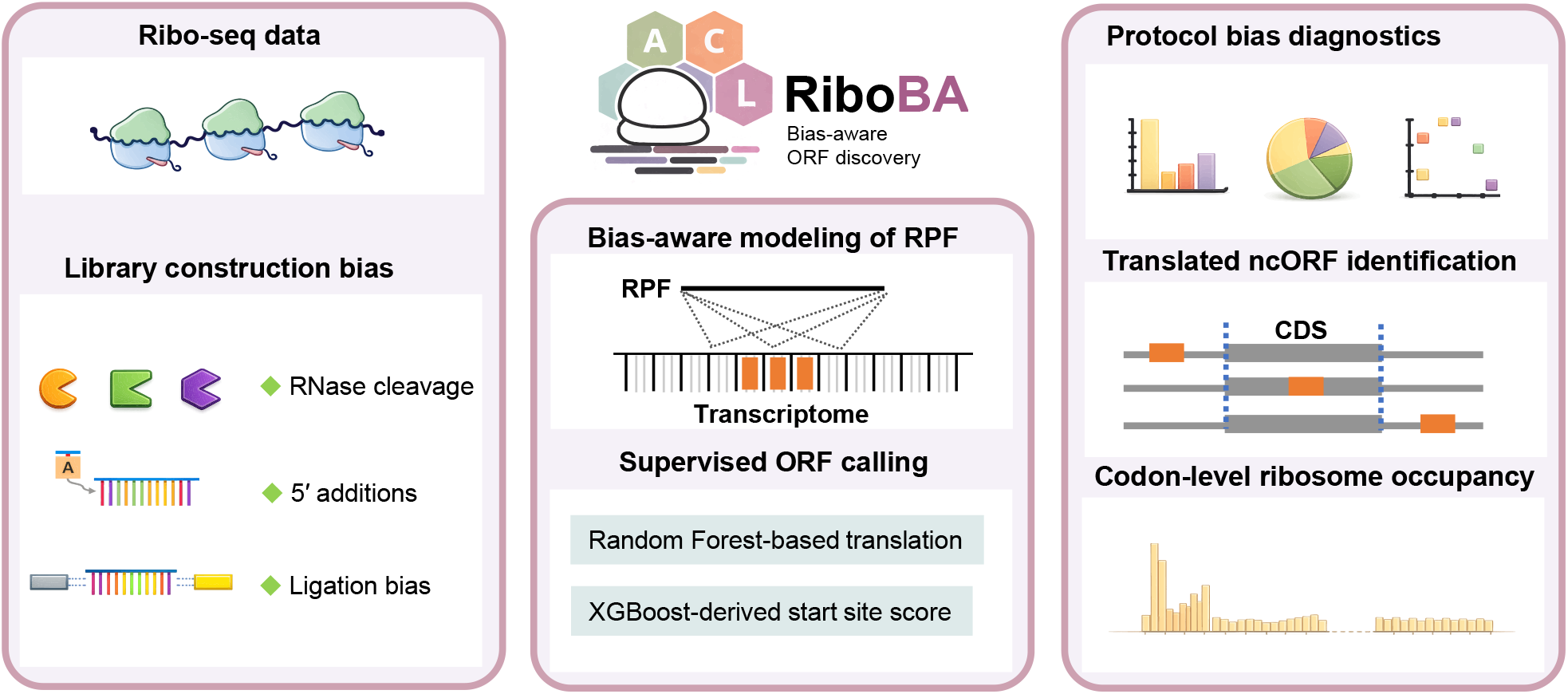

## Introduction

Ribosome profiling (Ribo-seq) captures genome-wide snapshots of translation by nuclease digestion of unprotected mRNA followed by deep sequencing of ribosome-protected fragments (RPFs) [1, 2]. Computational analysis of RPFs signals enables transcriptome-wide identification of actively translated open reading frames (ORFs) and has revealed extensive translation beyond canonical gene annotations [3, 4, 5]. These ncORFs arise in diverse genomic contexts [6]. Large-scale studies combining Ribo-seq with CRISPR-based screens have revealed that dozens to hundreds of ncORFs are essential for cell proliferation [5] and stress responses [7], and in some cases act through encoded microproteins involved in signaling, metabolic rewiring, and disease-related pathways [8]. In parallel, immunopeptidomic analyses have revealed that a substantial fraction of MHC class I–presented peptides originate from ncORFs, supporting their potential to generate antigenic peptides and contribute to immune recognition [9]. Despite their biological relevance [10], many ncORFs are short and often translated at relatively low levels [11], making them inherently difficult to identify with confidence [12].

Since the advent of Ribo-seq, considerable effort has therefore been devoted to developing computational methods for ncORF identification [13, 14]. These methods typically leverage characteristic signatures of translation inferred by mapping RPFs to the peptidyl (P)-site positions [13]. Computational approaches for identifying translated ncORF can be broadly categorized into three methodological classes: supervised classifiers, such as ORF-RATER [15] and RibORF [16], which apply regression or tree-based models to features reflecting periodicity and read coverage; (ii) generative probabilistic models, including PRICE [17] and Rp-Bp [18], which model codon-level translation activity using expectation–maximization or Bayesian inference; and (iii) periodicity-based statistical methods, such as RiboTaper [19], RiboCode [20], RiboTISH [21], and SPECtre [22], which directly assess 3-nt periodicity via spectral analysis or frame-specific statistical testing.

Although these methods differ in their downstream statistical frameworks, they generally share a common preprocessing strategy: P-site positions are assigned using fixed, length-specific offsets. This strategy introduces two major limitations. First, it typically restricts analysis to a subset of RPF lengths with strong periodicity and applies offset estimates that differ across tools [23], which may lead to signal loss and distortion. Second, simplifying RPFs to a fixed offset representation at the outset discards protocol-dependent information that could otherwise be modeled downstream. PRICE alleviates the first limitation by using probabilistic P-site assignment and retaining all RPF lengths, but it does not explicitly account for protocol-induced biases. A growing body of evidence demonstrates that systematic biases introduced during Ribo-seq library construction can profoundly distort RPF signals [24, 25, 26, 27]. Nuclease digestion bias introduces changes in RPF length and position: under-digestion produces extended overhangs that broaden the distribution of RPF lengths and obscure 3-nt periodicity [28], whereas over-digestion—particularly with RNase I—can dissociate ribosomal subunits and alter the distribution of RPFs [29]. Alternative nucleases such as MNase and P1 can reduce over-digestion [30, 31]. However MNase introduces cleavage bias [32], and the impact of these biases remains poorly addressed by existing ORF-calling methods. Ligation and circularization steps introduce additional biases, as enzymes such as T4 RNA ligase favor specific terminal nucleotides, resulting in a skewed representation of RPFs [33, 34, 35]. Furthermore, 5′ non-templated nucleotide additions introduced by reverse transcriptase cause systematic offset errors that disrupt 3-nt periodicity [36]. These biases may partly underlie the substantial discrepancies and limited reproducibility in ncORF identification reported across ORF-calling tools in recent benchmarking studies [37, 38].

To address these challenges, we developed RiboBA, a bias-aware probabilistic framework that explicitly models systematic biases in Ribo-seq data to improve ORF identification. RiboBA models footprint generation as a probabilistic mapping from latent P-site positions to observed RPFs, incorporating protocol-induced biases—including nuclease cleavage bias, the ribosome protection effect, ligation bias, and 5′ non-templated nucleotide additions—within a unified framework. The parameters associated with these protocol-induced biases are jointly inferred with codon-level ribosome occupancy using an expectation– maximization-like optimization strategy. RiboBA performs probabilistic P-site mapping by distributing each RPF across all geometrically compatible positions, weighting them by their posterior probabilities rather than assigning a single fixed site. By reweighting frame-ambiguous signals, this soft assignment corrects protocol-induced distortions and restores attenuated 3-nt periodicity. Leveraging adjusted P-site profiles, RiboBA computes translation-relevant features for both ORFs and initiation sites, which are evaluated via supervised learning models.

Using bias-aware simulations and analyses of human and *Drosophila* Ribo-seq datasets across multiple protocols, we show that RiboBA improves both canonical and non-canonical ORF detection compared to five state-of-the-art tools. (i) In simulations informed by RNase I, MNase, and P1-derived empirical parameters, RiboBA consistently achieved superior ORF detection performance. (ii) Across nine human datasets, RiboBA yielded highly reproducible ncORF detections with superior replicate concordance and validated immunopeptidomics evidence across multiple biotypes (iii) RiboBA robustly identifies conserved ncORFs with coding potential in *Drosophila melanogaster* MNase datasets, confirming its adaptability across distinct protocols. Despite its sophisticated modeling, RiboBA remains computationally efficient on large-scale datasets while providing bias metrics and ribosome occupancy profiles for downstream analysis.

## MATERIALS AND METHODS

### Reference construction and ORF search space

RiboBA constructs a transcript-level reference from the input genome and gene annotation. For each gene, RiboBA selects a single representative transcript using a deterministic rule that prioritizes the transcript with the longest coding sequence and resolves remaining ties by total transcript length. This strategy provides a consistent reference for ORF analysis while reducing ambiguity in read assignment caused by extensive exon sharing among transcript isoforms. Candidate ORFs are then exhaustively enumerated on each representative transcript by scanning for all candidate start codons and extending to the first in-frame stop codon (UAA, UAG, or UGA).

### Probabilistic modeling of RPF generation

RiboBA models RPF generation as a protocol-dependent transformation from latent ribosome P-site positions to observed reads. Aligned reads are collapsed into footprint classes *r* ≡(*s, ℓ*), defined by their 5′ terminal coordinate *s* and fragment length *ℓ*, with observed counts denoted as *y*_*r*_. To estimate protocol-induced bias parameters Θ, model fitting is using well-characterized, annotated coding sequences. Let *λ*_*p*_ represent the ribosome occupancy at candidate P-site *p*. The expected read count for class *r* is defined as a mixture over feasible P-sites:

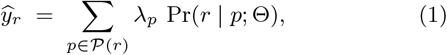

where 𝒫(*r*) is the set of P-sites compatible with class *r* given ribosome-geometry constraints on the 5′/3′ end-to-P-site distances for length *ℓ* (Supplementary Note S1).

The conditional probability Pr(*r* | *p*; Θ) is factorized as:

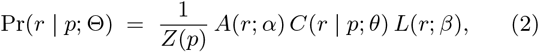

where *Z*(*p*) is the normalization constant over all footprint classes ℛ(*p*) compatible with P-site *p*. The factors *A, C*, and *L* represent 5′ additions, nuclease cleavage, and ligation efficiency, respectively.

Ribosome steric constraints define the admissible distances from P-site *p* to the 5′ and 3′ termini, denoted *d*_5_ and *d*_3_. Assuming conditional independence between termini given *p*:

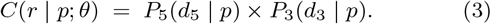

These probabilities capture both sequence preferences and distance-dependent protection effects. Let *x*_*t*_(*p, d*) be the nucleotide at distance *d* on terminus *t* ∈ {5, 3}, *b*_*x*_ the nucleotide-specific bias, and *h*_*t*_(*d*) the distance-dependent protection. The maintenance probability at distance *d* is defined as 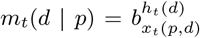. To model cleavage at distance *d*, we use a discrete hazard formulation over the admissible set 𝒟_*t*_:

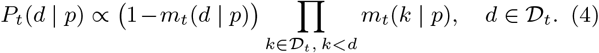

The normalization ensures 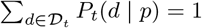.

Protocol-dependent 5′ additions, observed as first-base mismatches, are modeled by a library-specific categorical distribution *α* over *𝒜* = {*A, C, G, T*, ∅}, where ∅ denotes no addition.

To account for sequence-dependent fragment recovery, ligation efficiency is modeled as the product of terminus-specific probabilities:

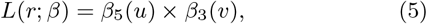

where *β*_5_(*u*) and *β*_3_(*v*) are ligation probabilities associated with 5′ and 3′ terminal *k*-mers *u* and *v* extracted from class *r*.

### Inference of model parameters and ribosome occupancies

Model parameters and P-site occupancies are inferred via a sequential procedure. First, the distribution of 5′ additions *α* is estimated by modeling mismatch compositions at the first aligned position relative to the reference. This approach recovers the full addition distribution, including cases where added bases coincide with the reference (Supplementary Note S2). The estimated *α* is then used to construct reweighted footprint counts 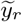.

We then infer P-site occupancies *λ*_*p*_ and cleavage-related parameters *θ* using an EM-like alternating optimization procedure. Let ℛ denote the set of retained footprint classes after filtering. For each *r* ∈ ℛ, the true P-site position *p* ∈ 𝒫(*r*) is treated as a latent variable. During this stage, the effects of ligation are held fixed, and we use Pr_0_(*r* | *p*; *θ*), the normalized conditional probability induced by the cleavage component(Eq. 3).

In the E-step, we compute the posterior probabilities over feasible P-sites:

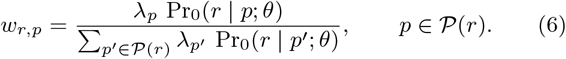

In the M-step, we update P-site occupancies by aggregating posterior-weighted reweighted counts:

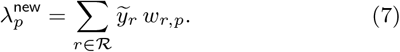

We update cleavage parameters *θ*, including sequence preferences at cleavage sites and distance-dependent steric protection effects, by maximizing the expected complete-data objective under *w*_*r,p*_ (Supplementary Note S3). The E- and M-steps are iterated until the parameter estimates converge.

After stabilizing (*λ, θ*), we estimate the ligation parameters *β* using the multiplicative terminal *k*-mer model (Eq. 5). We first compute the model-predicted baseline expectation 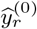 under the current estimates of (*λ, θ*) with ligation effects held fixed. We then estimate *β* using a regression model that treats the observed footprint-class counts as multiplicative deviations from 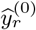. We use a log-link and include 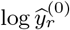 as an offset to isolate systematic deviations attributable to terminal ligation biases (Supplementary Note S4).

### Supervised identification of translated ORFs

Using bias-aware occupancies *λ* estimated in an all-frame manner, we performed supervised learning to identify translated regions and select the most probable initiation site within each.

Annotated CDS regions served as the primary positive training sets. To exclude weakly periodic or non-translating segments, we filtered these regions using a random forest classifier (`ranger` version 0.17.0, R) trained on well-supported CDS segments against an empirical null of randomly shifted read alignments. CDS segments passing this criterion formed the **long-ORF positive set**, while the **long-ORF negative set** combined these shift-based nulls with out-of-frame (+1 and −1) CDS controls. To address the scarcity of validated non-canonical translation events, such as short open reading frames (sORFs), we generated **pseudo-sORFs** from truncated long-ORF segments and **pseudo-overlap sORFs** by combining signals from overlapping structures.

To characterize these candidate regions, we extracted three feature classes: (i) *coverage and strength*, comprising the count and fraction of occupied codons and frame-stratified occupancy; *3-nt periodicity and frame consistency*, using frame usage proportions, log-odds for dominant-frame preference, and frame entropy; and (iii) *positional trends*, including 5′–3′ signal gradients and frame-consistency scores relative to empirical CDS distributions.

The `ranger` classifier was then trained on these features to assign translation probability scores. To account for heterogeneous background noise across ORF types, we applied class-specific decision thresholds. Candidates were categorized as CDS, uORF (upstream ORF), uoORF (upstream overlapping ORF), intORF (internal ORF), Ext (N-terminally extended CDS), Trunc (N-terminally truncated CDS), doORF (downstream overlapping ORF), dORF (downstream ORF), or Variant (novel transcript structures). Regions exceeding their respective thresholds were retained for initiation-site refinement.

To identify the most likely start codon within each translated region, we trained a binary XGBoost classifier (`xgboost` version 1.7.11.1, R). The training set utilized annotated CDS start sites as positives, with negatives derived from alternative in-frame ATGs located upstream or within the same CDS. Each candidate site was represented by (i) a 30-nt local P-site occupancy window and (ii) one-hot encoded flanking nucleotides. The decision threshold was optimized to maximize the F1 score on a held-out validation set. During inference, the first ATG exceeding this threshold was prioritized as the predicted initiation site; scoring of near-cognate start codons is also supported via user configuration.

Following ORF identification, we adjusted ribosome occupancy by restricting footprint assignment to in-frame positions within the detected ORFs. Finally, ORF coordinates were converted from transcript to genomic using annotated gene models. Each ORF was labeled by its position relative to annotated CDSs and flagged for in-frame overlap to facilitate downstream interpretation and visualization.

### Data processing and benchmarking

Human (HEK293/HEK293T) and *Drosophila* (embryo and S2 cell) Ribo–seq datasets were obtained from the NCBI SRA (Supplementary Table S1). Sequencing adapters were trimmed using Cutadapt (version 4.4), retaining 15–45 nt reads with base quality ≥20 to capture the full spectrum of RPFs generated under different experimental protocols. rRNA and tRNA reads were filtered by alignment to curated references using Bowtie (version 1.3.1) with up to two mismatches to sensitively remove contaminating reads. Remaining reads were aligned to the RiboBA-defined representative transcriptome (see Reference construction) using a two-pass strategy: reads were first mapped to annotated protein-coding transcripts, followed by alignment of unmapped reads to annotated long non-coding transcripts (Bowtie, up to one mismatch). This staged approach reduces spurious alignments to long non-coding transcripts.

To benchmark RiboBA, we selected representative tools from three methodological classes: supervised classifiers (ORF-RATER [15], RibORF [16]), generative models (PRICE [17]), and periodicity-based methods (RiboCode [20], RiboTISH [21]). Analyses used the hg38 (Ensembl version 109) and BDGP6.32 (Ensembl version 6.32.54) references for human and *Drosophila*, respectively. Candidate ORFs included both canonical (ATG) and near-cognate initiators. Tool-specific configurations and full commands are detailed in Supplementary Note S5 and Supplementary Code 1.

To enable cross-tool comparison, all predicted ORFs were converted to genomic coordinates and classified by their positions relative to annotated CDSs, with ORFs located on long non-coding transcripts designated as lncRNA-derived ORFs (lncORFs). Overlaps with annotated CDSs were assessed to determine frame-specific relationships; in-frame overlaps were excluded from downstream mass spectrometry validation to ensure the specificity of novel peptide identification.

### Simulation of ribosome profiling data

We simulated Ribo–seq libraries under three scenarios to evaluate parameter recovery and ORF detection (implementation details in Supplementary Code 1).

To evaluate parameter recovery, we simulated Ribo– seq libraries using the probabilistic mixture model defined in Eq. 1 under six protocol configurations: (i) RNase I (high dosage/high resolution), (ii) RNase I (low dosage/low resolution), (iii) MNase, (iv) nuclease P1, (v) RNase I with 5′ additions, and (vi) RNase I with sequence-dependent ligation bias. Codon-level occupancies *λ*_*p*_ were estimated from a representative HEK293T dataset (SRR23242344) and used as sampling probabilities in a multinomial distribution to simulate the distribution of ribosomes across P-site positions. For each configuration, we simulated 10 replicate libraries of 1×10^6^ reads each, using 1,000 randomly selected protein-coding genes.

To benchmark translation detection, we generated matched positive and negative sets through positional jittering of observed RPFs. Using RNase I (SRR23242345), MNase (SRR7073124), and nuclease P1 (SRR23242346) datasets, we selected 2,000 protein-coding genes with robust CDS-aligned read support. The positive set retained original RPF coordinates, while the negative set was constructed by jittering RPF positions within ±3 nt to disrupt 3-nt periodicity. The positive and negative RPF sets were combined with non-jittered reads from all other genes to form complete synthetic libraries, exported in FASTQ format with a constant Phred score of 40.

Finally, to assess ncORF detection, we simulated ncORF profiles by generating codon-level occupancies *λ*_*p*_ from CDS-derived donor distributions. These profiles were mapped to candidate ncORF coordinates and then used to synthesize read libraries under five protocol regimes: RNase I, MNase, nuclease P1, RNase I with 5′ additions, and RNase I with ligation bias. For each ncORF biotype, we generated a balanced set of 50 positive sets using bias parameters inferred from empirical datasets (SRR23242345, SRR7073124, SRR23242346). Negative sets were constructed using the same sampling procedure followed by nucleotide-scale P-site jittering to disrupt periodicity. These ncORF RPFs were combined with simulated CDS-derived background reads to form complete synthetic libraries, which were then exported in FASTQ format. RiboBA and alternative tools were evaluated using ROC and precision–recall (PR) curves via the `precrec` R package (version 0.14.0).

### Evolutionary conservation and coding potential analysis

Base-wise phyloP conservation scores [39] for *Drosophila melanogaster* (dm6, 27-way) were obtained from the UCSC Genome Browser. For each RiboBA-identified ORF, phyloP scores were extracted using `rtracklayer` (version 1.58.0), and the mean score was calculated as an ORF-level measure of evolutionary conservation.

To assess coding potential, PhyloCSF scores were extracted for ORFs ≥10 codons from the Broad Institute dm6 PhyloCSF track hub, which provides strand- and frame-specific scores for the dm6 assembly [40]. For each candidate, we retained the score corresponding to its identified strand and reading frame as a metric of protein-coding signature.

### Immunopeptidomics validation of identified ORFs

Evidence for the translation of identified ORFs was evaluated using public HLA-I immunopeptidomics data from HEK293 cells (PXD025449). To ensure specificity, in-frame ncORFs overlapping annotated CDSs were excluded. For each tool, identified ORFs were collapsed into a non-redundant set by grouping those sharing a stop codon and retaining only the most upstream start site.

These sequences were translated *in silico* to construct custom protein databases for each tool and Ribo–seq sample. HLA-I spectra were queried against these databases using `FragPipe` (version 22.0) and `MSFragger` (version 4.1) under a non-specific HLA workflow. False discovery rates (FDR) were controlled at 1% at both PSM and protein levels using `Philosopher`. Only uniquely mapping peptides (those assigned to a single ORF) were retained as evidence of translation.

## RESULTS

### RiboBA method overview

A schematic overview of RiboBA is shown in Fig. 1, with full technical details provided in the Materials and Methods and Supplementary Materials. In contrast to many existing ribosome profiling tools that assign P-site positions using fixed read–length-dependent offsets, RiboBA explicitly models the protocol-dependent process by which latent ribosomal P-sites give rise to observed RPFs. The generative model captures key technical biases introduced during library construction, including nuclease-specific cleavage bias, 5′-end non-templated nucleotide additions, and ligation efficiencies. Given the estimated parameters, RiboBA assigns each footprint a posterior over candidate P-sites based on its geometry. These weights are used to reassign frame-ambiguous signals, recovering P-site occupancy profiles with clear 3-nt periodicity. The resulting profiles are summarized into CDS-derived features that feed supervised machine-learning models for translated ORF detection. An EM-like alternating optimization scheme, combined with sparse matrix operations, accelerates P-site inference and feature extraction, enabling efficient analysis of large ribosome profiling data sets on standard workstations. Despite the increased model complexity, RiboBA does not impose prohibitive computational overhead, and in our benchmarks, it can achieve runtimes comparable to or faster than those of other widely used ORF-calling tools. RiboBA is implemented as an open-source R package available at https://github.com/Bai-JunYu/RiboBA.

**Figure 1.**
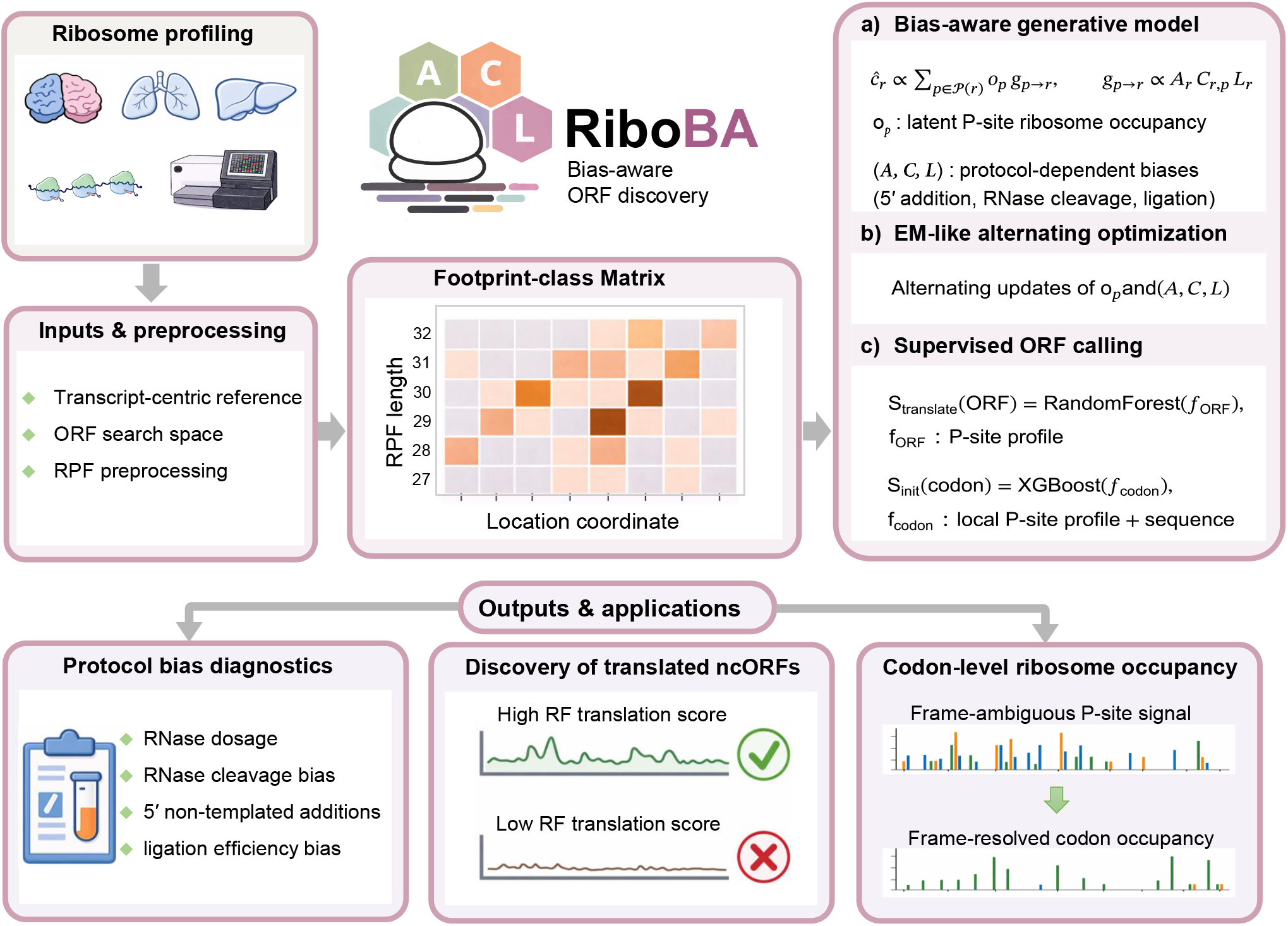
Schematic overview of the RiboBA framework. RiboBA is a bias-aware probabilistic framework for robust ORF identification from ribosome profiling data. The pipeline begins with preprocessing and aligning RPFs to a transcript-level reference to define the candidate ORF search space. A generative module then employs an EM-like optimization procedure to jointly infer protocol-induced biases, such as nuclease cleavage and ligation efficiency, alongside codon-level ribosome occupancy. These bias-adjusted profiles are subsequently utilized by a supervised module to score translated ORFs and initiation sites. The final outputs include protocol-bias diagnostics, identified ncORFs, and adjusted ribosome occupancy profiles for downstream functional analyses.

### Simulation-based recovery of protocol-induced biases

As ground truth for bias parameters is unavailable in empirical Ribo–seq data, we used our parameter-recovery simulations to assess the accuracy of RiboBA. We generated synthetic libraries across six protocol configurations (see Materials and Methods) using a fixed ribosome occupancy profile. Across all conditions, RiboBA accurately recovered the ground truth bias parameters (Supplementary Figure S1). Specifically, cleavage profiles for both termini were consistently inferred across RNase I (high- and low-dosage), MNase, and nuclease P1 regimes, with estimated probability curves closely matching the simulated truth (Supplementary Figure S1A).

Similarly, sequence-dependent ligation preferences were faithfully recovered: estimated 3-mer ligation efficiencies at the 5′ and 3′ termini showed strong Pearson correlations with the ground truth (average *r* ≈ 0.92; Supplementary Figure S1B– C). RiboBA also accurately inferred the base-specific cleavage preferences of MNase (Supplementary Figure S1D) and the probabilities of 5′ additions (Supplementary Figure S1E). Together, these results demonstrate that our generative model robustly infers protocol-induced bias parameters across diverse settings, providing a solid basis for subsequent bias-aware P-site inference and ncORF identification.

### Simulation-based benchmarking of ORF detection

We next evaluated whether RiboBA improves the identification of translated ORFs relative to existing methods using two simulation frameworks designed to assess detection accuracy under distinct protocol regimes.

The first benchmark, based on positional jittering, evaluated whether each method could distinguish translated CDSs from frame-disrupted signals, using a strategy similar to RiboCode [20]. Positive sets comprised 1,000 genes with CDS-aligned reads derived from empirical human Ribo–seq datasets (RNase I, MNase, or nuclease P1). Negative sets consisted of a separate set of 1,000 genes from the same datasets, where reads were subjected to positional jittering. This ±3-nt perturbation disrupts 3-nt periodicity while preserving the overall read distribution, providing a stringent benchmark for evaluating the translation specificity of identified ORFs. Across all three nuclease protocols, RiboBA consistently achieved the highest AUROC and AUPRC values (Supplementary Figure S2), outperforming RiboCode, PRICE, RibORF, and RiboTISH. ORF-RATER was excluded from this benchmark, as its supervised learning framework is not compatible with the structure of this simulation. These findings demonstrate that explicitly accounting for protocol-induced biases improves the robustness of CDS-level ORF identification under localized frame-disrupting perturbations.

The second benchmark, our synthetic ncORF detection framework, evaluated performance in identifying translated ncORFs under realistic protocol-induced biases. Positive sets comprised 250 translated ncORFs per protocol, simulated from CDS-derived P-site occupancy profiles and empirical bias parameters. Negative sets consisted of an equal number of non-translated ncORFs, generated using identical occupancy profiles but with positional jittering applied to P-site coordinates prior to footprint simulation. This approach ensures that 3-nt periodicity is disrupted while preserving global read distribution and protocol-induced biases, providing a rigorous test for the identification of non-canonical translation. We benchmarked each tool across simulated datasets spanning RNase I, P1, MNase, RNase I with 5′ additions, and RNase I with ligation bias. RiboBA consistently achieved the highest AUROC and AUPRC values (Figure 2A, Supplementary Figure S3A). Beyond accuracy, RiboBA exhibited the broadest coverage, effectively identifying nearly the entire positive and negative sets (Figure 2B, Supplementary Figure S3B). Among the alternative methods, PRICE achieved relatively high classification accuracy, second only to RiboBA, yet exhibited consistently limited coverage across protocols. RiboTISH provided comparatively broad coverage; however, its classification accuracy declined markedly under P1 and MNase regimes. RiboCode displayed a more balanced trade-off between accuracy and coverage overall, but experienced pronounced reductions in both metrics under MNase digestion. In contrast, RibORF and ORF-RATER showed consistently lower accuracy and more restricted coverage across all tested protocol conditions.

**Figure 2.**
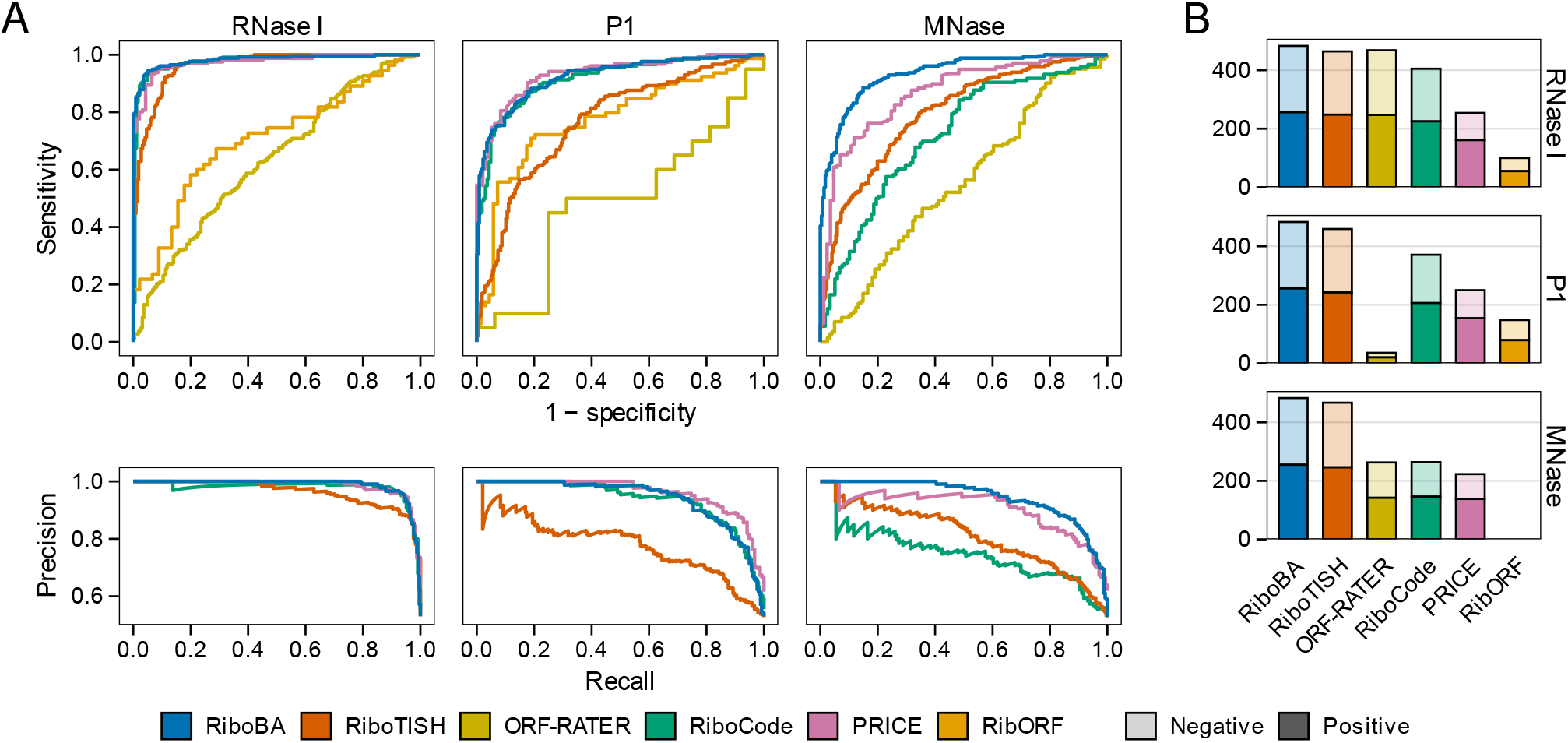
Benchmarking ncORF identification on simulated ribosome profiling datasets. Synthetic Ribo–seq datasets were generated under three nuclease conditions using empirical parameters from HEK293/HEK293T studies (RNase I and P1 from Lucas et al.; MNase from Darnell et al.). These datasets were used to evaluate the identification of translated ncORFs across diverse protocols. (A) Receiver operating characteristic (ROC) and precision–recall (PR) curves comparing RiboBA with PRICE, RiboCode, RibORF, RiboTISH, and ORF-RATER. Performance was assessed based on the method-specific identified ncORFs in each condition. (B) Total counts of identified positive and negative ncORFs across the three nuclease regimes.

This combination of high accuracy and broad detection coverage demonstrates that RiboBA provides a more versatile and generalizable framework for the identification of translated ncORFs across diverse experimental conditions. Despite its sophisticated modeling, RiboBA remains computationally efficient, with runtimes comparable to or faster than those of existing methods (Supplementary Figure S4).

### Protocol-induced bias in empirical datasets

We next evaluated whether the footprint-generation parameters inferred by RiboBA from empirical Ribo–seq datasets reflect biologically meaningful, protocol-induced biases. To enable cross-protocol comparisons while mitigating biological variability, we curated nine human HEK293/HEK293T Ribo– seq libraries representing diverse library construction protocols and known sources of technical bias.

Using this curated dataset, we first assessed the internal consistency of the protocol-induced parameters inferred by RiboBA within each library. Parameter estimates were independently computed from two random subsets of reads per library across ten replicate splits. The inferred ribosome protection profiles and 3-mer ligation efficiencies were highly consistent between splits, with Pearson correlation coefficients exceeding 0.99 (Supplementary Figure S5A-B). Similarly, MNase-specific base preferences and 5′ non-templated addition probabilities were highly reproducible (Supplementary Figure S5C-D), confirming the robustness of the inferred protocol-induced bias parameters.

With robust parameter inference confirmed across replicates, we next evaluated whether the inferred biases reflect biologically meaningful, protocol-induced features of ribosome footprints. We analyzed ribosome protection profiles flanking the P-site across nine protocols (Figure 3A). RNase I libraries generally exhibited narrow protection windows and steep gradients, consistent with shorter RPFs and more pronounced 3-nt periodicity. However, notable variability was observed across RNase I datasets. For instance, the Calviello library displayed sharp transitions in its protection profile, which were associated with strong periodicity. A similar trend was observed in the Martinez libraries: increasing nuclease concentrations resulted in progressively steeper protection profiles and stronger periodicity, further supporting this association. In contrast, the Lucas replicates yielded highly concordant protection profiles, underscoring the precision and biological fidelity of RiboBA-inferred parameters. Among non-RNase I protocols, MNase libraries showed broad, flat protection profiles, particularly at the 5′ end, consistent with prior findings that 3′-end-based offsets more reliably capture periodicity under such conditions [41]. P1 libraries exhibited intermediate steepness between RNase I and MNase, reflecting moderate 3-nt periodicity.

**Figure 3.**
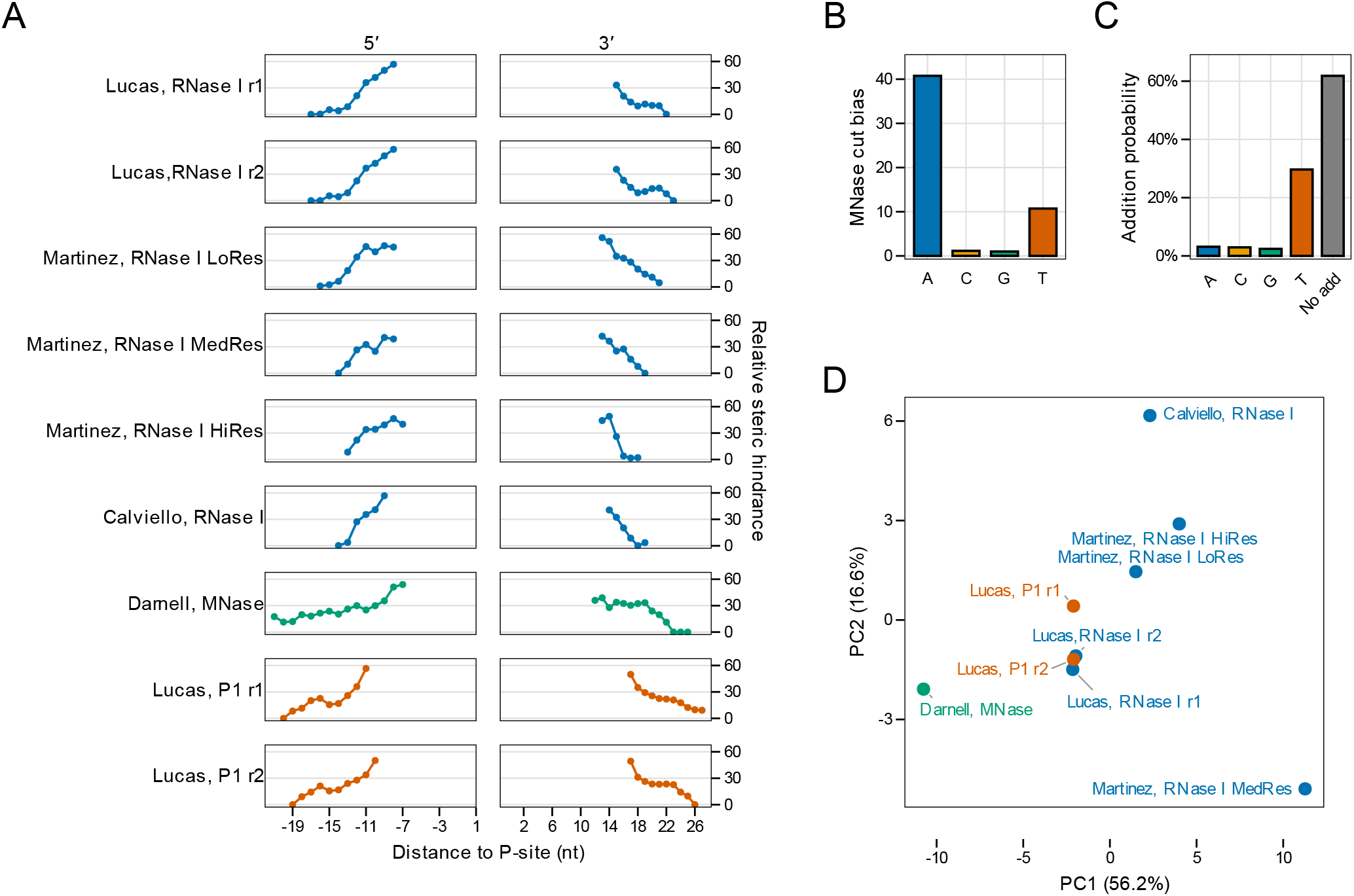
Protocol-induced bias parameters inferred by RiboBA from nine empirical HEK293/HEK293T Ribo–seq datasets. (A) Ribosome protection effects represented as terminus-specific relative steric-hindrance profiles flanking the P-site for 5′ and 3′ read ends; colors indicate the ribonuclease (RNase I, MNase, or P1), and higher values indicate increased resistance to nuclease cleavage. (B) MNase-specific cleavage bias in the Darnell dataset, shown as relative base cleavability across A/C/G/T, where larger values indicate higher cleavage susceptibility. (C) Estimated 5′ non-templated nucleotide addition probabilities for the Darnell dataset. (D) Principal component analysis (PCA) of inferred 3-mer ligation efficiencies at the 3′ end across the nine datasets; points represent individual libraries and are colored by ribonuclease type.

Next, we examined base-specific nuclease preferences. Biochemical studies have established that MNase preferentially cleaves A/T-rich regions over G/C-rich sites, with cleavage rates differing by up to 30-fold [32]. RiboBA accurately recapitulated this signature, inferring MNase cleavage rates approximately 40-fold and 10-fold higher for A and T, respectively, compared to C/G in the Darnell dataset (Figure 3B). In contrast, RNase I and P1 libraries exhibited minimal base-specific preferences (Supplementary Figure S6A), consistent with the known lower sequence specificity of these enzymes. These results demonstrate that RiboBA faithfully recovers nuclease-induced sequence biases from empirical Ribo–seq data.

We further examined 5′ non-templated nucleotide additions. RiboBA estimated low 5′-addition probabilities in most RNase I and P1 libraries, but detected markedly elevated levels in the Darnell MNase datasets (Figure 3C). To assess ligation efficiency bias, we summarized the inferred 3-mer ligation efficiencies at both termini and projected all libraries into a low-dimensional space via principal component analysis (PCA; Figure 3D; Supplementary Figure S6B-C). Samples from the same study clustered tightly, while remaining clearly separated from other datasets. This pattern indicates that the ligation parameters inferred by RiboBA capture study-specific signatures rather than stochastic technical noise. This pattern indicates that the inferred ligation parameters capture reproducible biochemical signatures—reflecting the inherent sequence preferences of ligases and adaptors—rather than stochastic technical noise.

Together, these results demonstrate that RiboBA enables precise and reproducible inference of protocol-induced biases across diverse empirical datasets. The inferred parameters may therefore serve as diagnostic metrics and design guides for optimizing future ribosome profiling experiments.

### Benchmarking ncORF identification on empirical Ribo–seq datasets

Having established that RiboBA accurately recovers protocol-induced biases, we next evaluated whether this modeling improves the identification of non-canonical ORFs in practice. We compared RiboBA with five widely used tools across nine human HEK293/HEK293T Ribo–seq libraries generated using RNase I, MNase, or nuclease P1 protocols.

We first assessed the ORF biotype composition identified by each method across the nine human libraries (Supplementary Figure S7). As expected, RNase I libraries—which typically exhibit stronger 3-nt periodicity—supported more translated canonical ORFs than MNase or P1 libraries. However, the degree of this protocol dependence varied significantly among tools (Supplementary Figure S7A). For the five alternative methods, translated canonical ORF counts fluctuated substantially across protocols, ranging from approximately 1k to 16k. This variability suggests that canonical ORF identification by these tools is strongly sensitive to protocol-dependent signal quality. In contrast, RiboBA yielded a more stable identification of 11k to 12k canonical ORFs across all protocols. An analogous pattern was observed for ncORFs (Supplementary Figure S7B), where RiboBA maintained robust identification of uORFs, uoORFs, intORFs, and lncORFs in MNase and P1 libraries, as well as in RNase I datasets with attenuated periodicity. Conversely, alternative methods frequently showed marked reductions under these biased or lower-signal conditions.

We next evaluated the reproducibility of ncORF identifications across biological replicates. Using the Lucas RNase I and P1 replicate pairs, we quantified concordance for each tool and ncORF biotype via both Jaccard similarity and the absolute number of shared identifications (Figure 4A). For uORFs and uoORFs, RiboBA achieved high replicate concordance while identifying substantially more candidates than alternative methods, a contrast that became more pronounced under the P1 protocol. For other biotypes, no single method consistently dominated, although ORF-RATER showed superior performance for intORFs in both shared identification count and replicate concordance.

**Figure 4.**
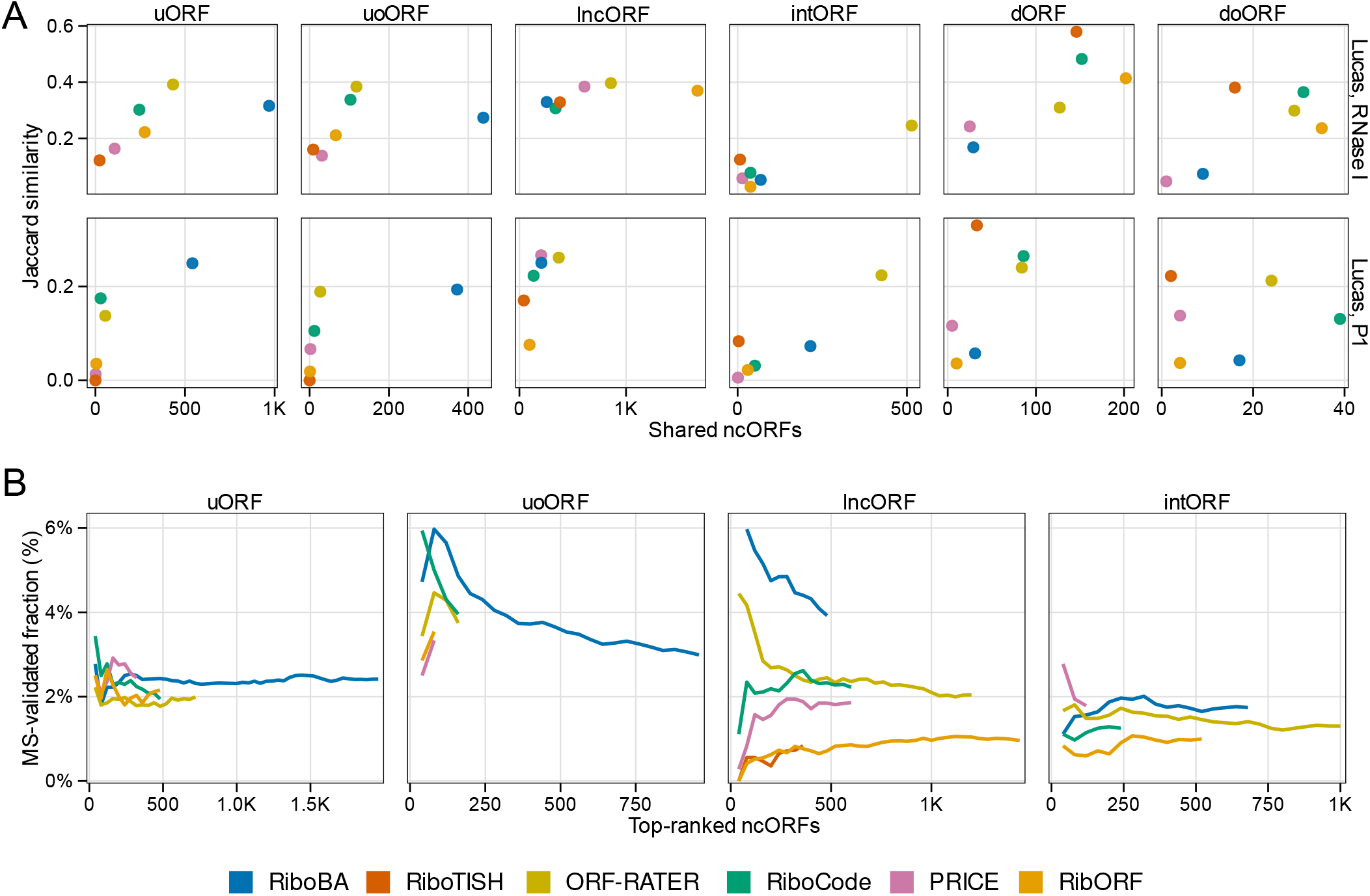
Reproducibility and immunopeptidomics validation of identified ncORFs. ncORFs were identified by RiboBA, PRICE, RiboCode, RibORF, RiboTISH, and ORF-RATER across nine empirical HEK293/HEK293T Ribo–seq datasets. (A) Jaccard similarity between biological replicate pairs (RNase I and P1 from Lucas et al.) versus the number of shared ncORF identifications, stratified by ncORF biotype. (B) HLA–I immunopeptidomics evaluation: identified ncORFs were ranked by method-specific scores, and the cumulative fraction of MS-validated ncORFs was plotted against the number of top-ranked identifications. Data are stratified by biotype (uORF, uoORF, intORF, and lncORF; dORF/doORF are omitted due to sparsity) and averaged across libraries (at least five datasets per data point).

Finally, we sought orthogonal evidence for identified ncORFs using HLA-I immunopeptidomics mass spectrometry— a stringent validation given the short length and low abundance of most ncORF-encoded microproteins [4]. By searching HEK293T MHC class I spectra against custom databases of identified ORFs, we quantified uniquely supported ncORFs after excluding in-frame CDS overlaps (Supplementary Figure S8). RiboBA yielded the highest numbers of MS-validated uORFs and uoORFs and ranked among the top methods for intORFs and lncORFs. In contrast, dORFs and doORFs were rarely detected by any method, a result consistent with the high prevalence of non-ribosomal protein-protected fragments in 3′ untranslated regions [12].

To evaluate score-based prioritization, we ranked identified ncORFs by their translation scores and assessed MS-support rates across score deciles (Figure 4B; dORFs and doORFs were excluded due to sparsity). Although overall validation rates hovered around 2%, distinct enrichment patterns emerged. For uORFs and intORFs, the frequency of MS support remained relatively stable across score ranks; nonetheless, RiboBA identified the largest cohort of MS-validated candidates while maintaining support rates comparable to or higher than those of alternative tools. Conversely, for uoORFs and lncORFs, high translation scores strongly correlated with MS-validated events. In these categories, top-ranked RiboBA identifications achieved validation rates of approximately 6%, outperforming all other methods in both absolute counts and support density for uoORFs. Notably, while RiboBA identified fewer lncORFs than ORF-RATER or RiboTISH, its hit rate was markedly superior (∼4% versus ∼1–2%), suggesting higher specificity in identifying bona fide non-canonical translation.

In summary, our multi-protocol evaluations and orthogonal immunopeptidomics evidence demonstrate the superior performance of RiboBA in identifying bona fide translation events. While alternative tools exhibit protocol-dependent fluctuations in sensitivity, RiboBA maintains stable and robust ncORF identifications even when 3-nt periodicity is attenuated, such as in MNase or P1 libraries. Beyond this robustness, RiboBA identifications of uORFs and uoORFs show higher reproducibility across biological replicates than existing methods. Furthermore, high-scoring RiboBA candidates are significantly enriched for MS-validated peptides, particularly within the uoORF and lncORF classes, where they achieve superior validation rates of approximately 6%. Collectively, these findings establish RiboBA as a reliable framework for ncORF identification that remains consistent across diverse experimental protocols.

### Case study: ncORF identification in Drosophila melanogaster

As a case study, we applied RiboBA to *Drosophila melanogaster*, a system inherently challenging for standard Ribo–seq protocols. Unlike human or yeast, *Drosophila* ribosomes possess an open architecture prone to RNase I-induced disassembly regardless of nuclease dosage [30]. This structural vulnerability necessitates alternative nucleases such as MNase to preserve ribosomal integrity; however, MNase introduces pronounced sequence-dependent cleavage biases that complicate ncORF identification [30]. To evaluate RiboBA’s robustness under these conditions, we analyzed three empirical MNase-based *Drosophila* datasets (Dunn, Greenblatt, and Zhang; Supplementary Table S1).

RiboBA consistently identified a robust set of ncORFs across the three *Drosophila* datasets. These candidates spanned diverse biotypes with reproducible distributions and stable initiation patterns (Supplementary Figure S9A–B). Although ATG was the predominant start codon, near-cognate initiators contributed a consistent fraction across all categories. Given that functionally relevant ncORFs often exhibit evolutionary constraints, we evaluated whether RiboBA identifications are enriched for conservation signals. We assessed these ncORFs against matched backgrounds using PhyloCSF for codon-level coding potential [40] and phyloP for nucleotide-level constraints [39]. Across all three datasets, RiboBA-identified ncORFs exhibited significantly elevated PhyloCSF scores relative to background, indicating stronger coding signatures (Figure 5A; Supplementary Figure S10A). Complementary phyloP analysis revealed ncORF subsets with nucleotide-level conservation comparable to annotated CDSs (Supplementary Figure S10B).

**Figure 5.**
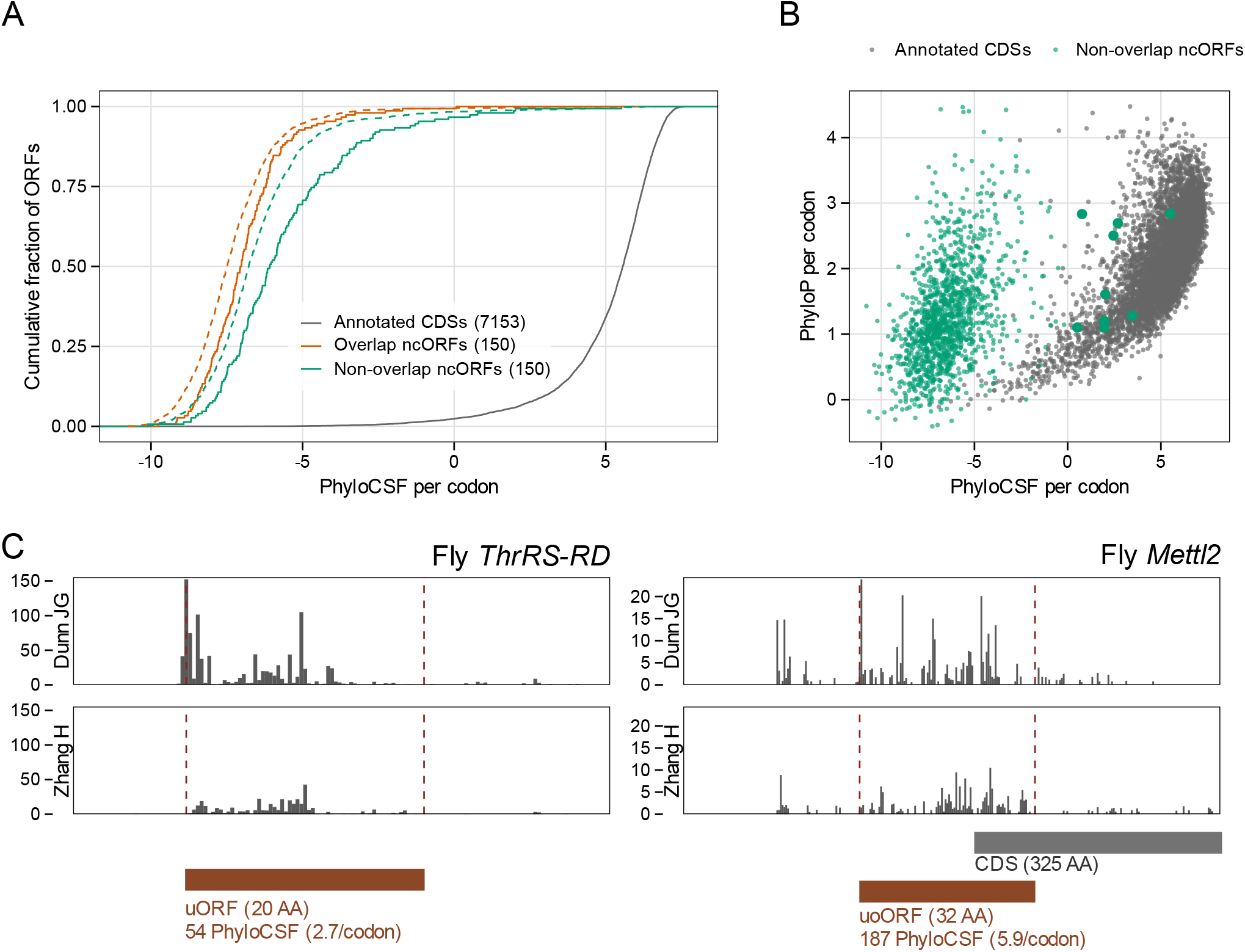
Codon-level conservation and occupancy profiles of RiboBA-identified *Drosophila melanogaster* ncORFs. (A) Empirical cumulative distribution functions (ECDFs) of per-codon PhyloCSF scores for annotated CDSs and RiboBA-identified ncORFs in the Zhang dataset. ncORFs are grouped into overlapping (uoORF, intORF, or doORF) and non-overlapping (uORF, dORF, or lncORF) categories. Dashed curves represent biotype-matched ORFs from the same transcripts that were not identified as translated. (B) Joint distribution of per-codon PhyloCSF and phyloP scores for annotated CDSs and non-overlapping ncORFs in the Zhang dataset. (C) P-site occupancy profiles for two conserved ncORFs identified across independent datasets (Dunn and Zhang): a uORF in *ThrRS* (ThrRS-RD) and a uoORF in *Mettl2*.

To jointly evaluate coding potential and nucleotide-level constraint, we analyzed both PhyloCSF and phyloP scores at the codon level and compared RiboBA-identified ncORFs with annotated CDSs (Figure 5B; Supplementary Figure S11). Annotated CDSs clustered distinctly within the high-PhyloCSF/high-phyloP domain, whereas only a small subset of ncORFs overlapped with this region, representing a conserved, coding-like minority. In contrast, a substantial fraction of identified ncORFs exhibited elevated phyloP but negative PhyloCSF scores. This pattern is consistent with conserved cis-regulatory roles rather than stable protein-coding functions, a trend particularly prominent among uORFs [10, 11, 42, 43].

Among the high-confidence ncORFs with positive PhyloCSF scores (Supplementary Table S3), RiboBA identified a conserved, translated uORF in *ThrRS* (FBgn0027081) and a uoORF within *Mettl2* (FBgn0035247) across independent datasets (Figure 5C). Both genes play essential roles in threonine-linked decoding: *ThrRS* encodes the threonyl-tRNA synthetase for charging tRNA^Thr^, which also functions as a non-canonical translation initiation factor [44], while *Mettl2* encodes a conserved tRNA methyltransferase that, in mammals, installs m^3^C at position 32 of tRNA^Thr^ isoacceptors [45]. The recurrent identification of upstream translation in both a synthetase and its corresponding tRNA modifier suggests a potential regulatory axis coordinating threonine-specific translational fidelity. In summary, our analysis of *Drosophila* datasets demonstrates that RiboBA effectively overcomes protocol-specific enzymatic biases to identify biologically relevant and evolutionarily conserved ncORFs. These findings highlight the capacity of RiboBA to uncover novel translational control mechanisms in experimentally challenging systems.

## DISCUSSION

Systematic biases introduced during Ribo–seq library preparation frequently obscure the 3-nt periodicity required for the precise identification of translated ncORFs [46]. To address this, we developed RiboBA, a bias-aware generative framework that models footprint generation as a probabilistic mapping from latent P-site positions to observed reads, followed by supervised learning to score candidate ORFs based on bias-adjusted ribosome occupancy profiles. Through extensive benchmarking using both simulations and multi-protocol empirical datasets from human and *Drosophila*, RiboBA consistently demonstrated superior accuracy and reproducibility compared with existing state-of-the-art tools. Collectively, our findings establish RiboBA as a robust, bias-aware solution that enables consistent ncORF identification across diverse experimental regimes.

The bias profiles inferred by RiboBA represent more than auxiliary parameters for ORF identification; they serve as interpretable diagnostic readouts to inform experimental protocol optimization. For example, the inferred sharpness of ribosome protection provides a quantitative metric for nuclease digestion stringency, enabling researchers to evaluate whether the applied nuclease dosage is excessive or insufficient. Other bias components capture distortions introduced at specific stages of library construction, thereby pinpointing technical steps that distort RPF signals. Beyond individual experiments, these inferred bias profiles collectively define sample-specific bias signatures—standardized representations of protocol-induced bias across datasets. Such signatures provide a foundation for the large-scale integration of ribosome profiling data from multiple laboratories. As the field moves toward unified and comprehensive translatome annotation [47, 48], RiboBA facilitates cross-study integration by explicitly accounting for dataset-specific systematic characteristics in heterogeneous libraries, thereby supporting more reliable normalization and comparison across studies.

Despite these advances, the identification of short and lowly expressed ncORFs remains a fundamental challenge. While RiboBA substantially improves sensitivity, increasing sequencing depth alone cannot fully compensate for the intrinsic limitations of RPFs, which are short and often unevenly distributed along transcripts. Furthermore, our current implementation adopts a simplified one-representative-transcript-per-gene strategy. In reality, translation is far more complex, frequently involving multiple isoforms with overlapping sequences that lead to substantial alignment ambiguity [49, 50]. To address these limitations, future extensions of RiboBA will focus on integrating signals across multiple Ribo–seq datasets. By transitioning from single-library modeling to an integrative, cross-protocol framework, we aim to enhance the high-resolution annotation of the dark translatome.

## Abbreviations

RPF: ribosome-protected fragment
Ribo-seq: ribosome profiling sequencing
ORF: open reading frame
ncORF: non-canonical ORF
RNase I: ribonuclease I
MNase: micrococcal nuclease
P1: nuclease P1

## Conflicts of interest

The authors declare that they have no competing interests.

## Funding

This work was supported by the Fund of Northwest A&F University (Z111021404), the “100-Talent Program” of Shaanxi Province of China (A289021612), and the National Natural Science Foundation of China (Grant No. 32270608) to R.Y.

## Data availability

The RiboBA package is available at https://github.com/Bai-JunYu/RiboBA. Detailed step-by-step instructions for annotation construction, data preprocessing, and package usage are provided in the repository documentation. All scripts used for benchmarking, downstream validation, and figure generation are available in the companion repository RiboBA-analysis at https://github.com/Bai-JunYu/RiboBA_analysis. Figure-ready processed data used to generate the main and supplementary figures have been deposited in Zenodo under DOI: https://doi.org/10.5281/zenodo.19070763.

## Author contributions statement

J.B. conceived the study, developed the RiboBA framework, performed all analyses, and drafted the manuscript. R.Y. supervised the project and revised the manuscript. Both authors read and approved the final manuscript.

## Acknowledgments

The authors thank members of the Yang laboratory for helpful discussions.

## Notes

https://doi.org/10.5281/zenodo.19070763

https://github.com/Bai-JunYu/RiboBA

https://github.com/Bai-JunYu/RiboBA_analysis

### Competing Interest Statement

The authors have declared no competing interest.

