## Supplementary Information for "RiboBA: a bias-aware probabilistic framework for robust ORF identification across diverse ribosome profiling protocols"

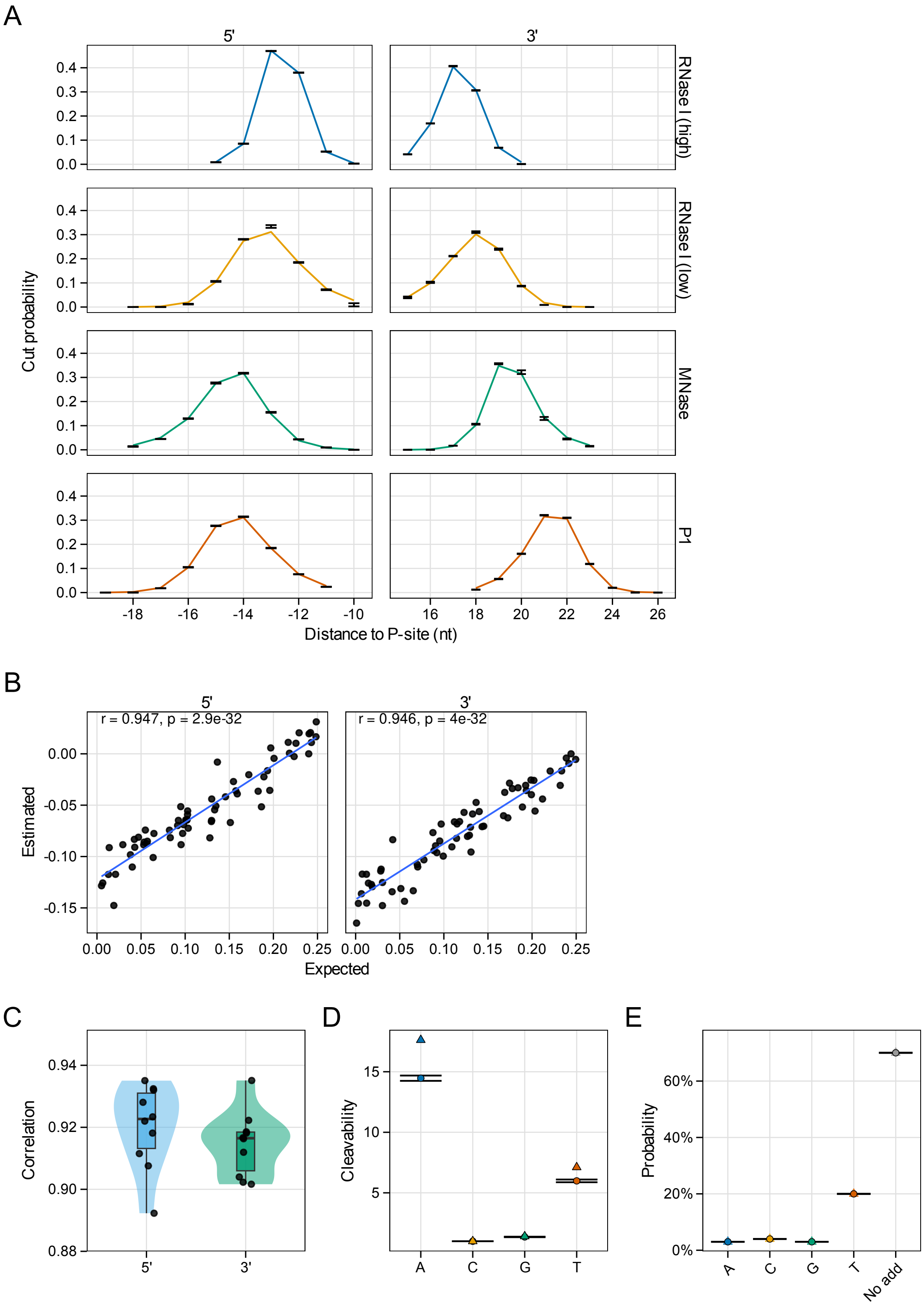
A
5'
3'
RNase I (high)
RNase I (low)
MNase
P1
Cut probability
Distance to P-site (nt)
B
r = 0.947, p = 2.9e-32
r = 0.946, p = 4e-32
Estimated
Expected
C
Correlation
D
Cleavability
E
Probability
No add

**Supplementary Figure S1. Recovery of protocol-induced bias parameters in simulations. (A)** Ground-truth versus RiboBA-estimated terminus-specific cut probability profiles flanking the P-site for RNase I (high/low dosage), MNase, and P1 (mean $\pm$ SD; $n = 10$), shown separately for 5′ and 3′ read ends. **(B–C)** Recovery of ligation bias parameters. **(B)** Correlation between mean estimated and expected 3-mer ligation efficiencies across 10 simulations, with Pearson coefficients indicated for both termini. **(C)** Pearson correlations between estimated and ground-truth ligation parameters across 10 independent simulated libraries. **(D)** MNase-specific cleavage bias shown as relative cleavability across A/C/G/T (triangles, expected; circles, estimated; mean $\pm$ SD; $n = 10$). **(E)** 5′ non-templated nucleotide addition probabilities (triangles, expected; circles, estimated; mean $\pm$ SD; $n = 10$).

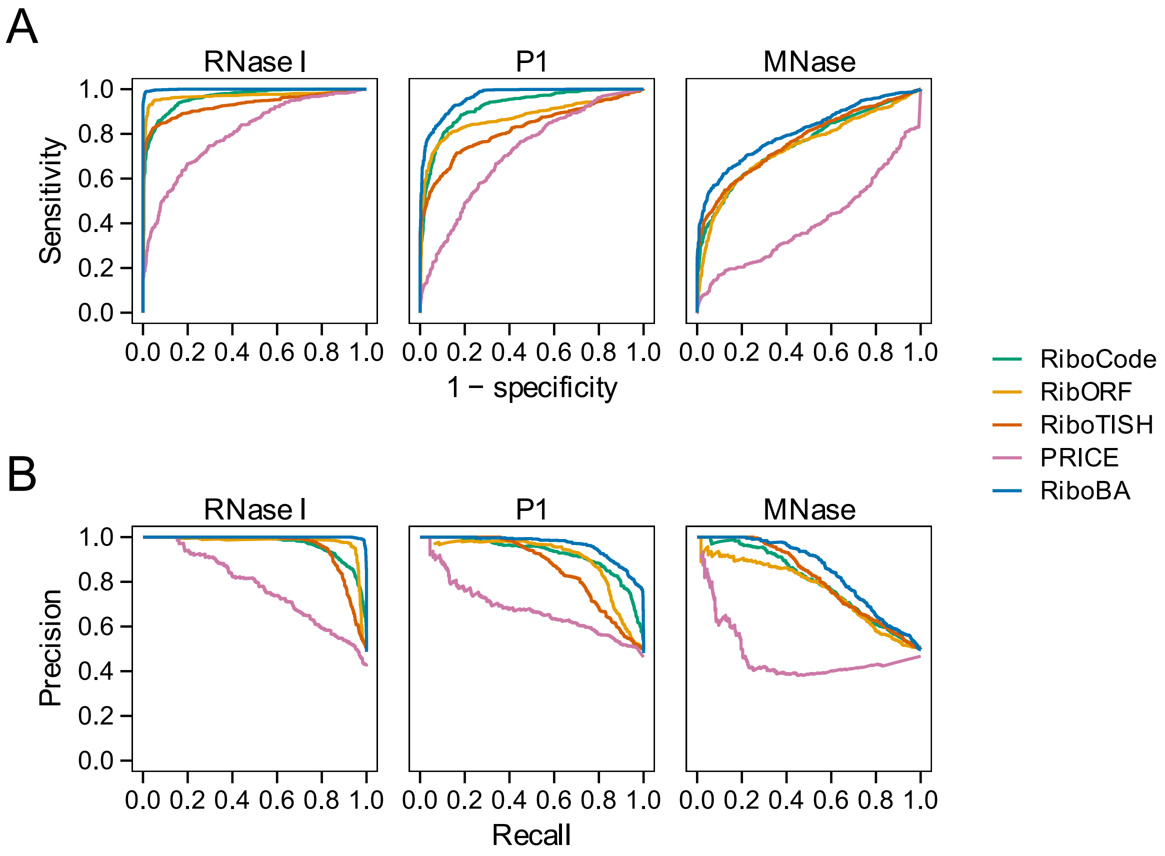


**Supplementary Figure S2. Performance benchmarking on CDS-level Ribo–seq simulations.** Synthetic datasets were derived from empirical HEK293/HEK293T Ribo–seq experiments (RNase I and P1 from Lucas *et al.*; MNase from Darnell *et al.*). **(A)** Receiver operating characteristic (ROC) curves comparing RiboBA with PRICE, RiboCode, RibORF, and RiboTISH. **(B)** Precision–recall (PR) curves comparing the same five tools. ORF-RATER was excluded from this benchmark because its supervised scoring framework is incompatible with the positional-jittering simulation design.

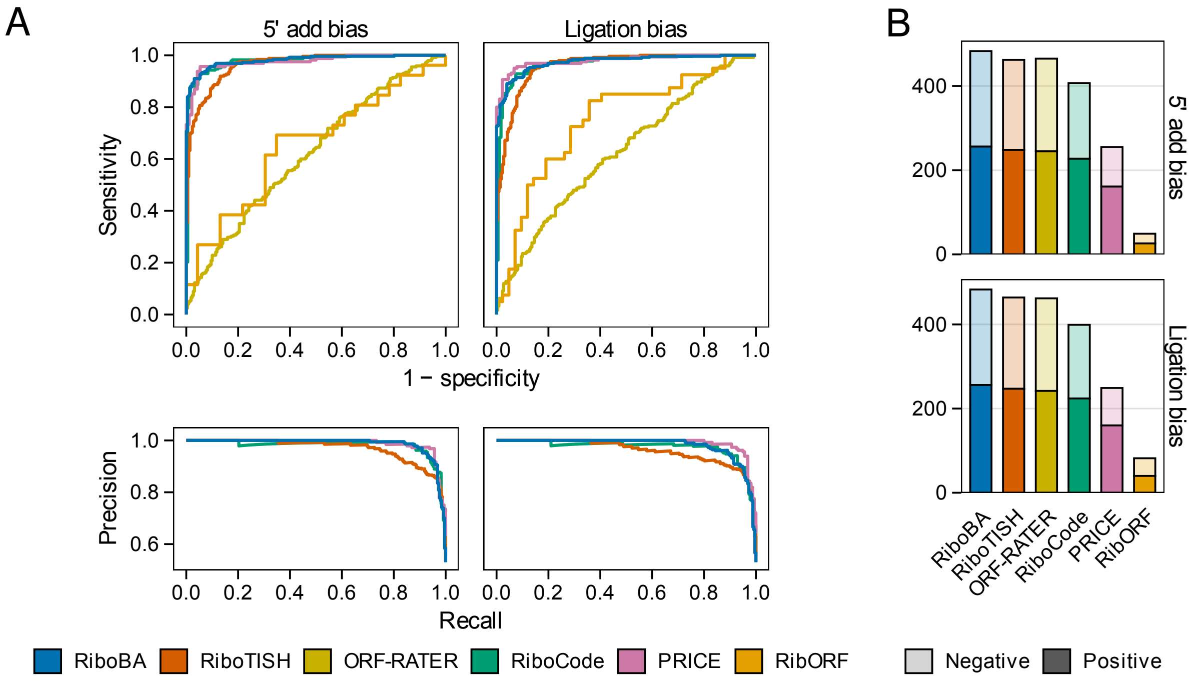


**Supplementary Figure S3. Performance of ncORF identification under simulated protocol-induced biases.** The benchmarking analysis was performed on synthetic datasets incorporating 5′ non-templated addition or ligation bias (parameters derived from empirical RNase I data, Lucas *et al.*). **(A)** Receiver operating characteristic (ROC) and precision–recall (PR) curves computed from identified positive and negative ncORFs for each method. **(B)** Total counts of identified positive and negative ncORFs across each protocol configuration.

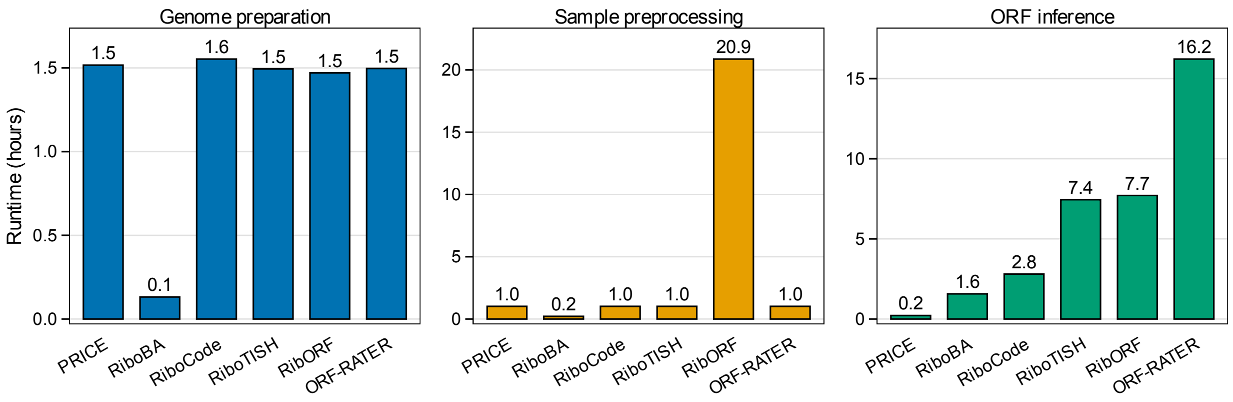


**Supplementary Figure S4. Computational efficiency of ncORF identification.** Total CPU time (core-hours) required for the identification of translated ncORFs from the Lucas RNase I human HEK293T Ribo–seq dataset, partitioned into genome preparation, sample preprocessing, and ORF-level inference.

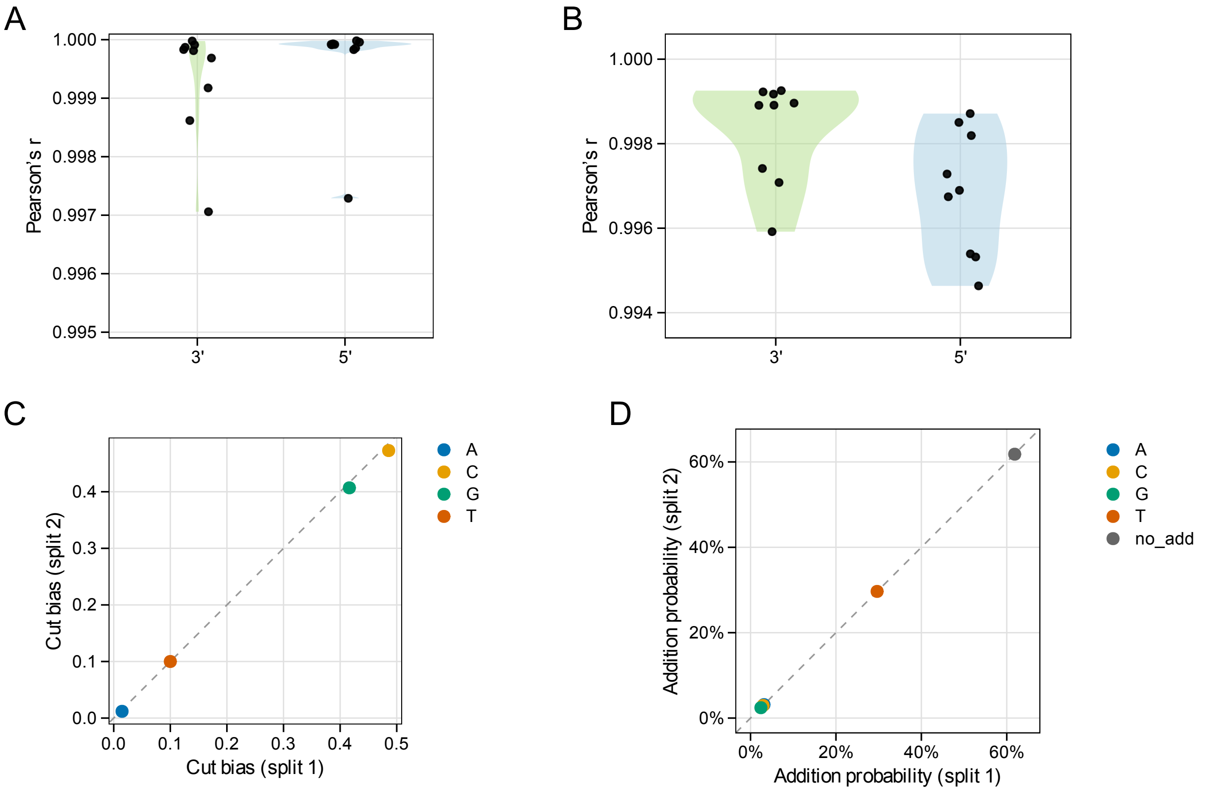


**Supplementary Figure S5. Internal consistency of protocol-induced bias parameters inferred by RiboBA from empirical HEK293/HEK293T Ribo–seq datasets. (A–B)** Pearson correlation coefficients ($r$) between two random read partitions for ribosome protection effects **(A)** and 3-mer ligation efficiencies **(B)**, evaluated for 5′ and 3′ termini (each point represents one dataset). **(C–D)** Agreement between partitions for MNase cleavage bias **(C)** and 5′ non-templated addition probabilities **(D)** in the Darnell dataset. In scatter plots, the dashed line indicates $y = x$.

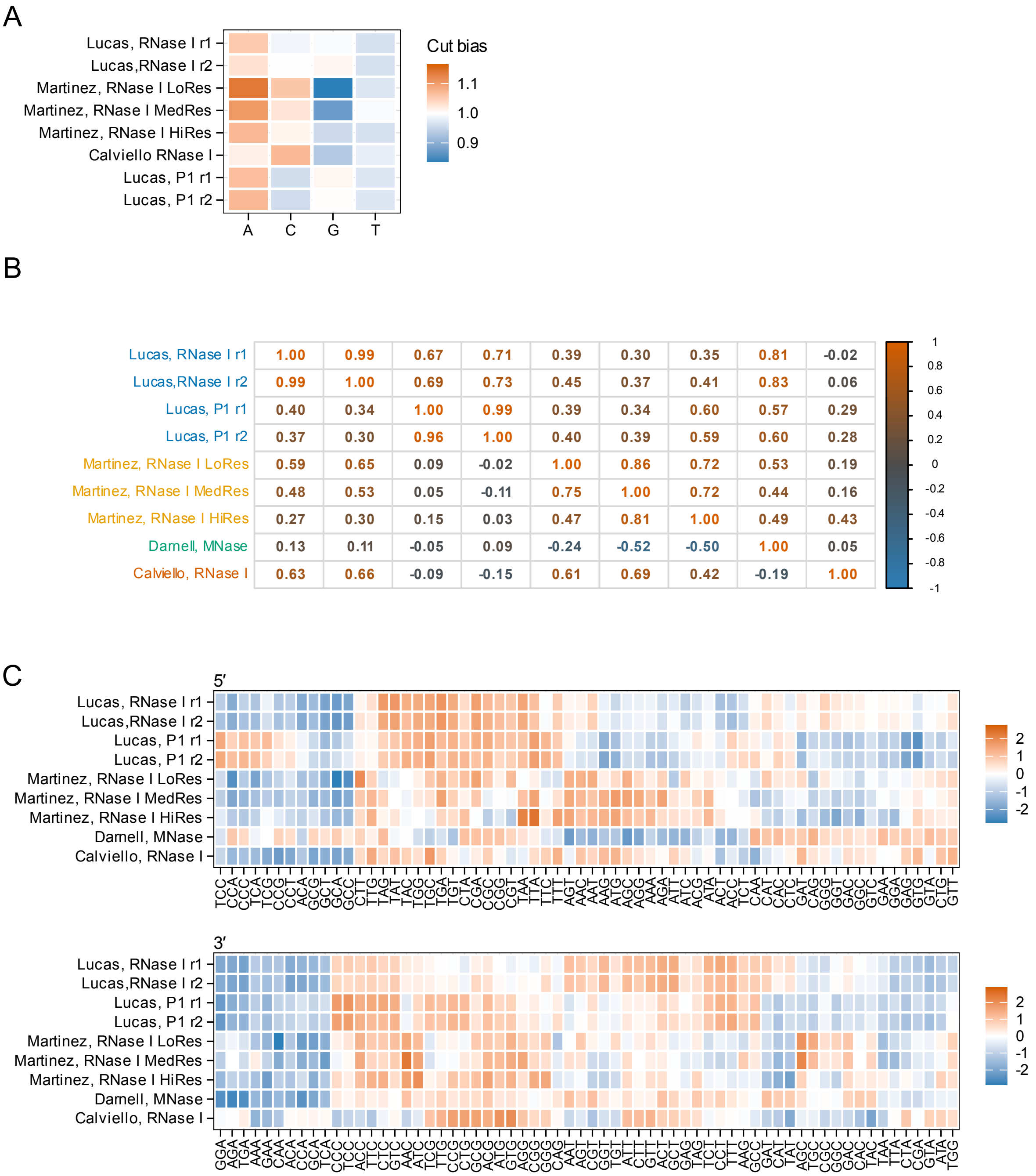
A
Lucas, RNase I r1
Lucas,RNase I r2
Martinez, RNase I LoRes
Martinez, RNase I MedRes
Martinez, RNase I HiRes
Calviello RNase I
Lucas, P1 r1
Lucas, P1 r2
A C G T
Cut bias
1.1
1.0
0.9
B
Lucas, RNase I r1 1.00 0.99 0.67 0.71 0.39 0.30 0.35 0.81 -0.02
Lucas,RNase I r2 0.99 1.00 0.69 0.73 0.45 0.37 0.41 0.83 0.06
Lucas, P1 r1 0.40 0.34 1.00 0.99 0.39 0.34 0.60 0.57 0.29
Lucas, P1 r2 0.37 0.30 0.96 1.00 0.40 0.39 0.59 0.60 0.28
Martinez, RNase I LoRes 0.59 0.65 0.09 -0.02 1.00 0.86 0.72 0.53 0.19
Martinez, RNase I MedRes 0.48 0.53 0.05 -0.11 0.75 1.00 0.72 0.44 0.16
Martinez, RNase I HiRes 0.27 0.30 0.15 0.03 0.47 0.81 1.00 0.49 0.43
Darnell, MNase 0.13 0.11 -0.05 0.09 -0.24 -0.52 -0.50 1.00 0.05
Calviello, RNase I 0.63 0.66 -0.09 -0.15 0.61 0.69 0.42 -0.19 1.00
1 0.8 0.6 0.4 0.2 0 -0.2 -0.4 -0.6 -0.8 -1
C
5′
Lucas, RNase I r1
Lucas,RNase I r2
Lucas, P1 r1
Lucas, P1 r2
Martinez, RNase I LoRes
Martinez, RNase I MedRes
Martinez, RNase I HiRes
Darnell, MNase
Calviello, RNase I
TCC CCA CCC TCA TCG CCG CCT ACA GCG GCT GCA GCC CTT TTG TAG TAT TAC TGG TGC TGA TGT CTA CGA CGC CGG CGT TAA TTA TTC TTT AGT AAC AAT AAG ATG AGC AGG AAA AGA ATT ATC ACG ATA ACT ACC TCT CAA CAT CAC CTC GAT CAG GGG GGT GAC GGC GTC GAA GGA GAG GTG GTA CTG GTT
2
1
0
-1
-2
3′
Lucas, RNase I r1
Lucas,RNase I r2
Lucas, P1 r1
Lucas, P1 r2
Martinez, RNase I LoRes
Martinez, RNase I MedRes
Martinez, RNase I HiRes
Darnell, MNase
Calviello, RNase I
GGA AGA TGA AAA GAA CAA ACA CCA GCA TCA CCC TCC ACC TTC CTC GTC AAC ATC TCG TTG CCG CTG GCG ACG ATG AGG GGG CAG AAT AGT CGT GGT TGT ATT CTT GTT ACT GCT GAG TAG TCT CCT TTT AAG GCC GAT CAT TAT AGC TGC CGC GGC GAC CAC TAC TAA CTA CGA GTA ATA TGG
2
1
0
-1
-2

**Supplementary Figure S6. Characterization of nuclease cleavage and ligation biases across nine empirical HEK293/HEK293T Ribo–seq datasets. (A)** Base-specific nuclease cleavage bias across A/C/G/T shown as relative cleavability for the eight non-MNase datasets. **(B)** Pearson correlation matrix of inferred 3-mer ligation efficiencies across datasets (5′ terminus, lower triangle; 3′ terminus, upper triangle). **(C)** Heatmaps of inferred 3-mer ligation efficiencies for 5′ (top) and 3′ (bottom) termini across all libraries.

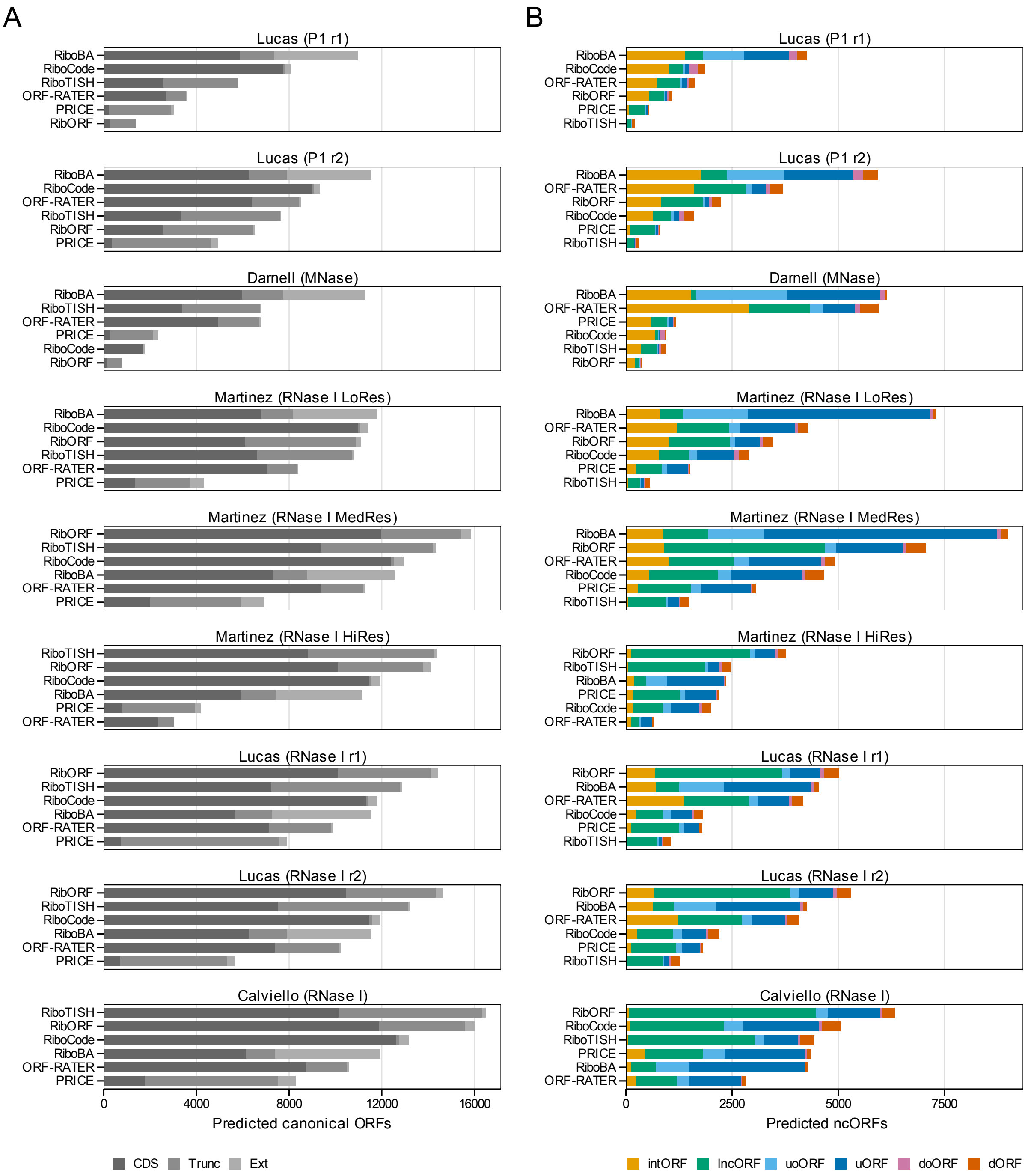

A
B
Lucas (P1 r1)
Lucas (P1 r2)
Darnell (MNase)
Martinez (RNase I LoRes)
Martinez (RNase I MedRes)
Martinez (RNase I HiRes)
Lucas (RNase I r1)
Lucas (RNase I r2)
Calviello (RNase I)
RiboBA
RiboCode
RiboTISH
ORF-RATER
PRICE
RibORF
0
4000
8000
12000
16000
Predicted canonical ORFs
CDS
Trunc
Ext
2500
5000
7500
Predicted ncORFs
intORF
lncORF
uoORF
uORF
doORF
dORF

**Supplementary Figure S7. Biotype composition of identified ORFs across nine empirical human Ribo–seq datasets.** ORFs identified by each method were quantified across nine human Ribo–seq datasets and grouped by biotype. **(A)** Canonical ORFs, including annotated, truncated, and extended CDS regions. **(B)** Non-canonical ORFs (ncORFs), including uORF, uoORF, intORF, dORF, doORF, and lncORFs.

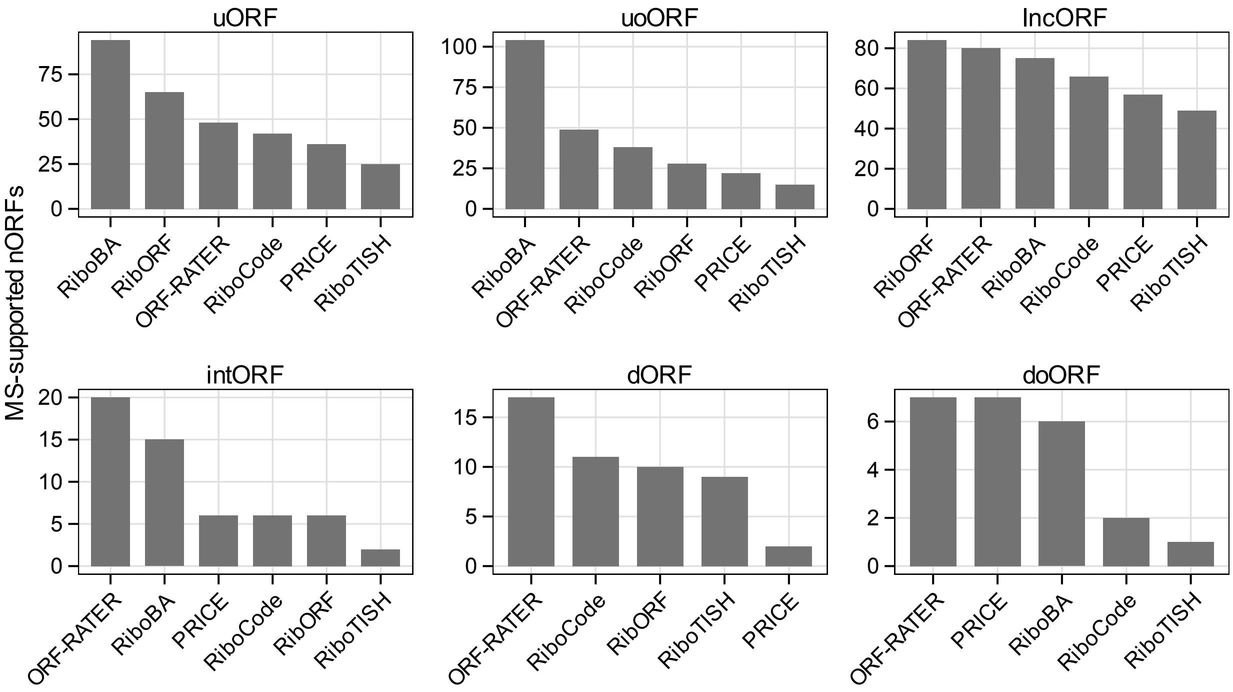


**Supplementary Figure S8. MS validation of identified ncORFs across nine empirical HEK293/HEK293T Ribo–seq datasets.** Identified ncORFs were evaluated using independent HLA–I immunopeptidomics mass spectrometry evidence. Bars represent the number of uniquely MS-validated ncORFs for each method, stratified by ncORF biotype.

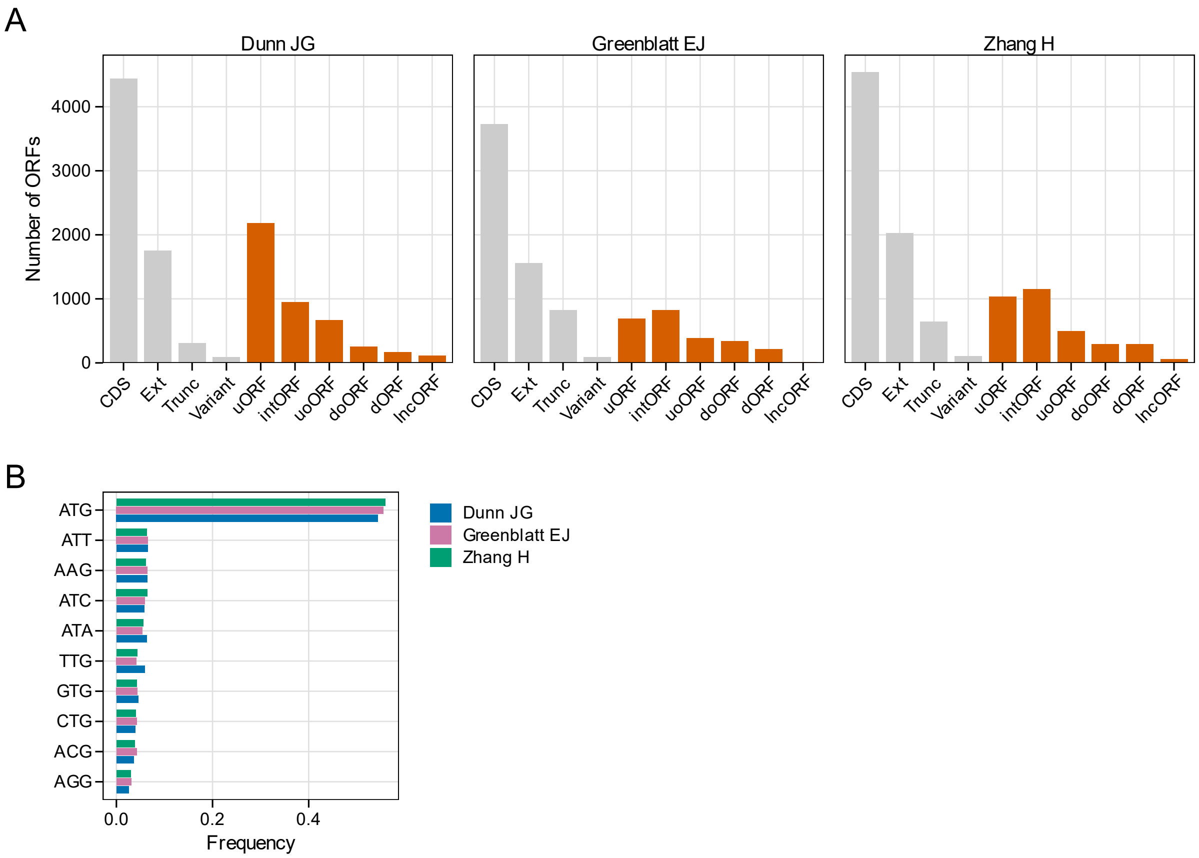


**Supplementary Figure S9. Summary of identified ORFs in empirical *Drosophila* Ribo–seq datasets.** Identified ORFs were compiled from three *Drosophila melanogaster* Ribo–seq datasets. **(A)** Total counts of identified ORFs by biotype in each dataset (with ncORF biotypes highlighted). **(B)** Start-codon usage across datasets, shown as the frequency of each initiator among identified ncORFs.

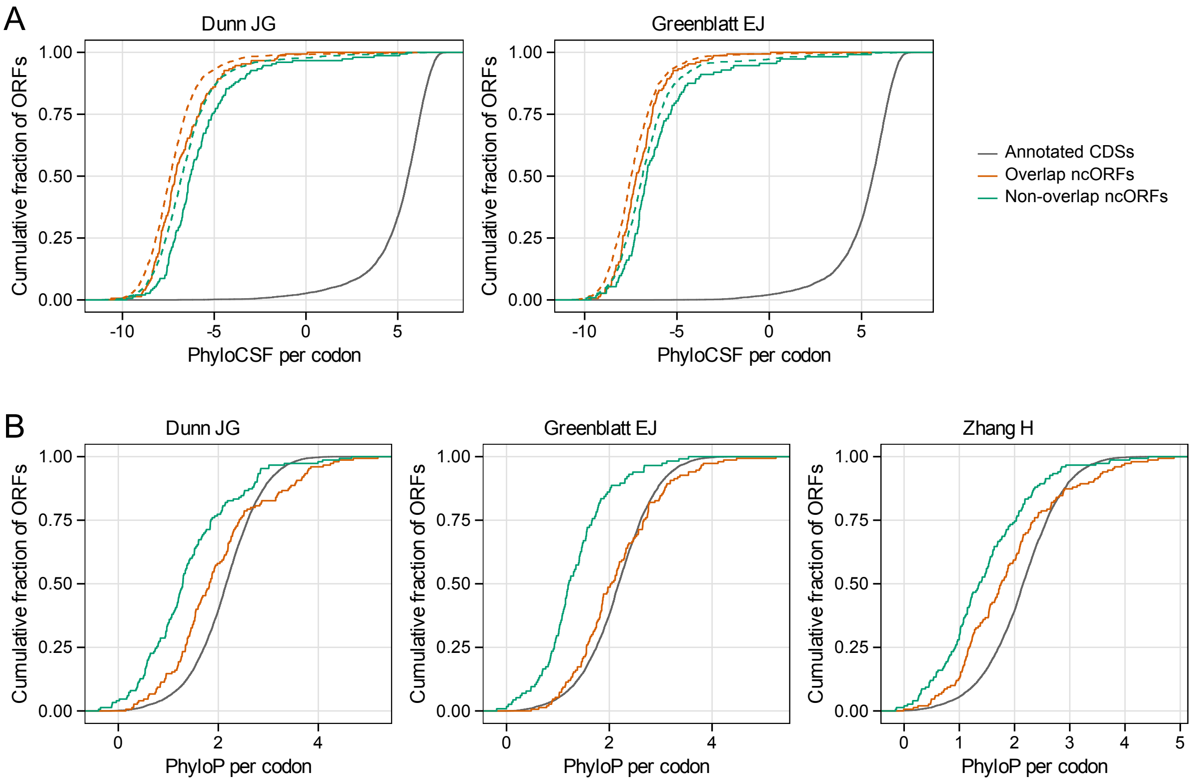


**Supplementary Figure S10. PhyloCSF and phyloP distributions of RiboBA-identified *Drosophila* ncORFs.** Empirical cumulative distribution functions (ECDFs) of per-codon PhyloCSF and phyloP scores are shown for annotated CDSs and top-ranked ncORFs (top 50 per biotype ranked by footprint support). ncORFs are grouped into overlapping (uoORF, intORF, or doORF) and non-overlapping (uORF, dORF, or lncORF) categories. **(A)** PhyloCSF ECDFs for the Dunn and Greenblatt datasets; dashed curves represent biotype-matched sequences from the same transcripts that were not identified as translated. **(B)** phyloP ECDFs for the Dunn, Greenblatt, and Zhang datasets.

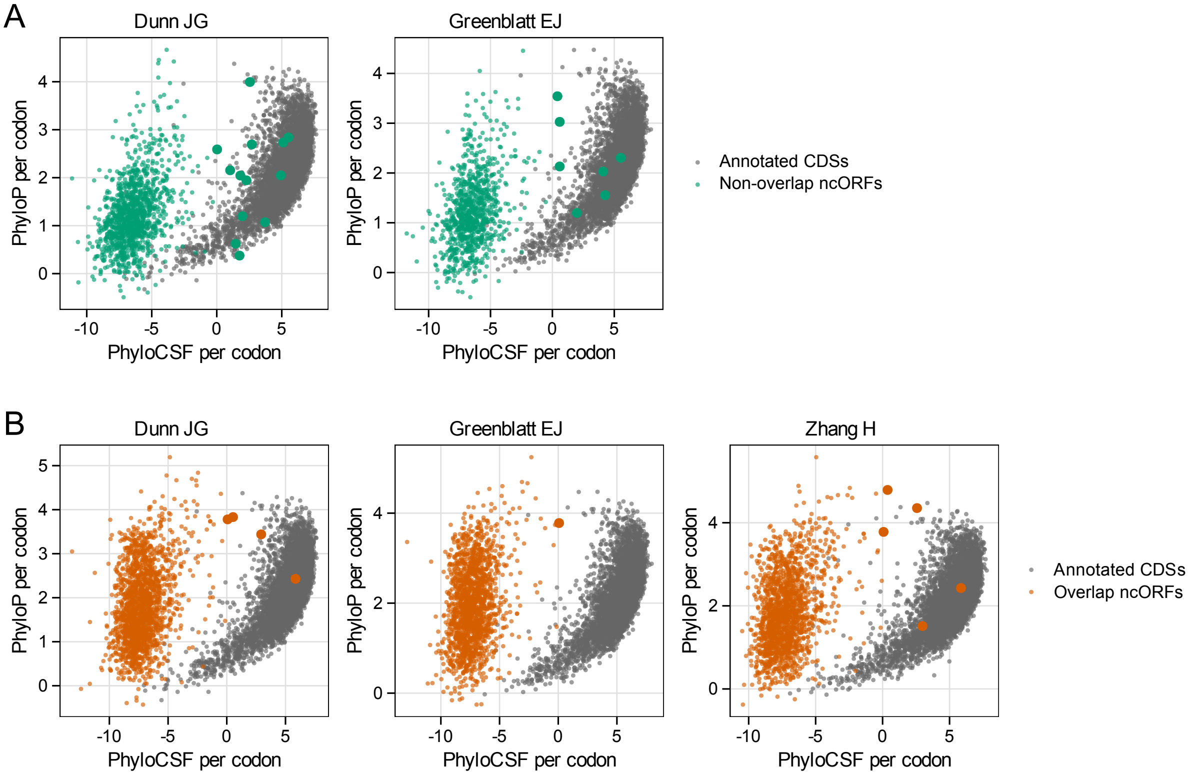


**Supplementary Figure S11. Joint distribution of PhyloCSF and phyloP scores for RiboBA-identified *Drosophila* ncORFs.** Scatter plots of per-codon PhyloCSF versus phyloP scores for annotated CDSs and identified ncORFs in empirical *Drosophila* Ribo–seq datasets. **(A–B)** ncORFs are shown separately for **(A)** non-overlapping (dORF, lncORF, and uORF) and **(B)** overlapping (doORF, intORF, and uoORF) categories. Points represent the top 1,000 identified ncORFs per biotype ranked by footprint support; ncORFs with positive PhyloCSF scores (PhyloCSF $>$ 0) are highlighted.

### Supplementary Note

#### 1. Ribosome-geometry constraints and P-site initialization

RiboBA employs a discrete ribosome footprint layout parameterized by $(L_5, S_5, S_3, L_3)$, where $L_5$ and $L_3$ specify the admissible distance ranges between the P-site and the 5′ and 3′ termini, respectively, and $S_5$ and $S_3$ define protected offsets. We initialize these parameters based on the minimum retained footprint length and the observed length distribution. To optimize the layout, we optionally apply an iterative adjustment heuristic that shifts boundary values by one nucleotide when the inferred distance-dependent steric-hindrance profiles concentrate probability mass at the most distal admissible distances.

Given the footprint layout parameters $(L_5, S_5, S_3, L_3)$, each footprint class $r \equiv (s, \ell)$ defines a feasible set $\mathcal{P}(r)$ of candidate P-site positions. A position $p$ is included in $\mathcal{P}(r)$ if it satisfies the discrete end-to-P-site distance constraints for length $\ell$, subject to two biological constraints: $p$ must be in-frame with and contained within the annotated CDS. These feasible sets $\mathcal{P}(r)$ specify the candidate P-site positions for the mixture model in Eq. (1) and define the support over which the conditional distribution $\Pr(r \mid p; \Theta)$ in Eq. (2) is normalized.

Specifically, $\delta_\ell$ is determined by aggregating the 5′ termini of reads of length $\ell$ into a metagene profile around annotated start codons. Within an admissible range (typically 12–15 nt), $\delta_\ell$ is defined as the distance that maximizes the read count, effectively representing the most frequent offset for that fragment length. For each footprint class $r \equiv (s, \ell)$, we then assign an initial P-site position as:

$$p = s + \delta_\ell,$$

which provides the baseline ribosome occupancy profile for initializing the probabilistic model.

#### 2. Estimating the 5′ non-templated addition distribution

We model 5′ non-templated additions using a library-specific distribution $\alpha = (\alpha_A, \alpha_C, \alpha_G, \alpha_T, \alpha_\varnothing)$, representing the probabilities of adding one of the four nucleotides or no addition ($\alpha_\varnothing$). Parameter estimation is based on three assumptions: (i) first-base mismatches at the 5′ terminus of addition-eligible reads arise predominantly from non-templated additions rather than sequencing errors or alignment artifacts; (ii) the composition of added bases is independent of the reference nucleotide at the first aligned position; and (iii) additions are restricted to at most one extra nucleotide.

Let $x \in \{A, C, G, T\}$ denote the reference nucleotide at the first aligned position and $a \in \{A, C, G, T\}$ denote the first sequenced base of the read. For non-templated addition-eligible reads exhibiting a mismatch ($x \neq a$), we summarize the observed counts in a mismatch table $M_{x,a}$. We first estimate the conditional distribution of added nucleotides given an addition event, denoted by $\tilde{\alpha} = (\tilde{\alpha}_A, \tilde{\alpha}_C, \tilde{\alpha}_G, \tilde{\alpha}_T)$, where $\tilde{\alpha}_a \geq 0$ and $\sum_a \tilde{\alpha}_a = 1$. The expected probability $p_{x \to a}(\tilde{\alpha})$ of observing base $a$ as a mismatch at reference position $x$, conditional on an addition event, is defined as:

$$p_{x \to a}(\tilde{\alpha}) = \frac{\tilde{\alpha}_a}{1 - \tilde{\alpha}_x}, \qquad a \in \{A, C, G, T\} \setminus \{x\}.$$

The corresponding observed mismatch fractions are calculated as:

$$\widehat{p}_{x \to a} = \frac{M_{x,a}}{\sum_{b \neq x} M_{x,b}}.$$

We then obtain the estimate $\widehat{\tilde{\alpha}}$ by minimizing the squared difference between the expected and observed fractions on the log scale:

$$\widehat{\tilde{\alpha}} = \arg\min_{\tilde{\alpha}} \sum_x \sum_{a \neq x} \left[ \log \widehat{p}_{x \to a} - \log \left( \frac{\tilde{\alpha}_a}{1 - \tilde{\alpha}_x} \right) \right]^2,$$

subject to $\tilde{\alpha}_a \geq 0$ and $\sum_a \tilde{\alpha}_a = 1$.

In the second step, we recover the overall addition rate. Let $\pi_x$ denote the background frequency of reference nucleotide $x$ among the mapped 5′ positions of addition-eligible reads, and let $m$ denote the observed fraction of such reads exhibiting a mismatch at the first aligned position. Under this model, the mismatch fraction satisfies:

$$m = \pi \Big( 1 - \sum_x \pi_x \tilde{\alpha}_x \Big),$$

where $\pi$ represents the probability of a non-templated addition event. This relationship yields the estimator:

$$\widehat{\pi} = \frac{m}{1 - \sum_x \pi_x \widehat{\tilde{\alpha}}_x}.$$

We then combine the estimated overall addition rate $\widehat{\pi}$ with the conditional composition $\widehat{\tilde{\alpha}}$

to obtain the five-category addition distribution $\widehat{\alpha}$:

$$\widehat{\alpha}_{\varnothing} = 1 - \widehat{\pi}, \qquad \widehat{\alpha}_a = \widehat{\pi}\,\widehat{\widetilde{\alpha}}_a, \quad a \in \{A, C, G, T\}.$$

The distribution $\widehat{\alpha}$ is used to correct 5′ terminus positions prior to EM-like fitting by redistributing reads across footprint classes. For reads with a first-base mismatch, the model deterministically attributes the mismatch to a non-templated addition, and their contribution is shifted to the corrected footprint class. For reads where the first base matches the reference, the presence of a non-templated addition is treated as a latent variable. Using the estimated model, we compute the posterior probability of a 5′ addition for each read and partition its contribution between the original and shifted footprint classes, ensuring that the total count $\sum_r \widetilde{y}_r = \sum_r y_r$ is preserved.

#### 3. Inferring cleavage-related parameters $\theta$

With $\lambda$ fixed, we update $\theta$ by matching the observed footprint-class counts with model-predicted expectations. Let $\widehat{y}_r(\theta)$ denote the expected count for footprint class $r$ under the generative model defined in Eq. (1), evaluated using current parameters; when decoupling ligation effects, the ligation term $L(r;\beta)$ is set to 1. To stabilize fitting across varying count magnitudes, we compare the observed counts $y_r$ and transformed expectations $\widehat{y}^*_r(\theta)$, where:

$$\widehat{y}^*_r(\theta) = k_r \widehat{y}_r(\theta),$$

and $k_r$ is an optional multiplicative factor that incorporates estimated terminal ligation efficiencies (e.g., $k_r = L(r;\widehat{\beta})$). We then minimize a robust discrepancy between these observed and expected counts:

$$\widehat{\theta} = \arg\min_{\theta \in \Omega_\theta} \sum_{r \in \mathcal{R}} \frac{\left|\widehat{y}^*_r(\theta) - y_r\right|}{\widehat{y}^*_r(\theta) + \left(\widehat{y}^*_r(\theta)\right)^2/10}, \tag{1}$$

where $\Omega_\theta$ imposes box constraints on the nuclease-bias parameters ($10^{-6} \le b_x \le 1 - 10^{-6}$ and $10^{-8} \le h_t(d) \le 1 - 10^{-6}$ for all nucleotides $x$ and distances $d$). The objective function in Eq. (1) is optimized using constrained derivative-free numerical routines under $\Omega_\theta$.

#### 4. Estimation of ligation parameters

Following the stabilization of $(\lambda, \theta)$, we attribute the remaining systematic discrepancies between observed footprint counts and baseline expectations to sequence-dependent ligation efficiency. Let $\widehat{y}_r^{(0)}$ denote the baseline expected count for footprint class $r$, computed under current $(\lambda, \theta)$ with ligation effects held constant (i.e., $L(r; \beta) = 1$). We then employ a regression model with a log-link and a structural offset:

$$\log(y_r) = \log(\widehat{y}_r^{(0)}) + \eta_0 + \eta_5\big(k_5(r)\big) + \eta_3\big(k_3(r)\big) + \varepsilon_r,$$

where $k_5(r)$ and $k_3(r)$ represent the 5′ and 3′ terminal $k$-mers of footprint class $r$, $\eta_0$ is a global intercept, and $\eta_5(\cdot)$ and $\eta_3(\cdot)$ denote additive terminal $k$-mer effects on the log scale. The fitted values define a multiplicative correction factor:

$$L_{\text{reg}}(r) = \exp\Big\{\eta_0 + \eta_5\big(k_5(r)\big) + \eta_3\big(k_3(r)\big)\Big\}$$

This factor is mapped to the multiplicative terminal $k$-mer model described in the Main Text by setting:

$$\beta_5(u) \propto \exp\{\eta_5(u)\}, \qquad \beta_3(v) \propto \exp\{\eta_3(v)\},$$

where normalizing constants are chosen such that $0 < \beta_5(u) < 1$ and $0 < \beta_3(v) < 1$ for all terminal $k$-mers $u$ and $v$. To ensure robust estimation for rare terminal contexts, the regression is performed on a balanced subset of footprint classes.

#### 5. ORF identification using alternative tools

The specific parameters and pipelines used for each tool are summarized below; detailed scripts and configuration files for all benchmarking analyses are provided at `https://github.com/Bai-JunYu/ORF-calling`.

RibORF (version 2.0) was executed on TopHat2 (version 2.1.1) alignments generated using Bowtie2 (version 2.5.1) with a maximum of two mismatches. Candidate ORFs with a minimum length of 6 codons (including both ATG and non-ATG starts) were identified using `ORFannotate.pl`. P-site offsets were inferred via `readDist.pl` and refined with `offsetCorrect.pl`. Final translated ORFs were identified per sample using `ribORF.pl`.

RiboCode (version 1.0.0) utilized a reference transcriptome constructed from Ensembl GTF and GRCh38 FASTA files via `prepare_transcripts`. Reads were aligned using

STAR (version 2.7.10b) in end-to-end mode ($\leq$2 mismatches) with unique mapping enforced. P-site offsets and valid read lengths (19–42 nt; $f_0 \geq 0.4$) were inferred from metagene plots, and translated ORFs were subsequently identified.

PRICE, as implemented in GEDI (version 1.0.6), utilized STAR (version 2.7.10b) to generate coordinate-sorted BAM files, allowing up to 2 mismatches and 20 multimapping alignments. Following genome indexing, ORFs were identified using the `Price` module, and outputs were converted from CIT to BED format using `ViewCIT`.

ORF-RATER was applied to the coordinate-sorted BAM files used for PRICE. Transcript annotations were converted to BED12 format and redundant isoforms were pruned. Candidate ORFs were generated using `find_orfs`, followed by P-site inference via `psite_trimmed.py`. Translation was modeled using `regress_orfs.py` and quantified from the resulting BED outputs.

RiboTISH (version 0.2.7) was executed on the same BAM files as PRICE. Footprint periodicity and valid read lengths (19–42 nt) were assessed using a periodicity threshold of 0.4. ORFs were identified using `ribotish predict` with the `--framebest` option to prioritize the most periodic frame; the `--alt` flag was enabled to allow for non-ATG initiation.
